# Extracellular vesicle cargo metabolome changes in response to the mesenchymal stromal cell microenvironment and influences cell quiescence and activation in a human breast cancer cell model

**DOI:** 10.1101/2022.12.16.520731

**Authors:** Sara Bartlome, Yinbo Xiao, Ewan Ross, Matthew John Dalby, Catherine Cecilia Berry

## Abstract

Breast cancer is the leading cause of cancer mortality in women worldwide and commonly metastasizes to the bone marrow, drastically reducing patient prognosis and survival. In the bone marrow niche, metastatic cells can enter into a dormant state, thereby evading immune surveillance and treatment, and can be reactivated to enter a proliferative state due to poorly understood cues. Mesenchymal stromal cells (MSCs) maintain cells in this niche partly by secreting extracellular matrix and paracrine factors and by responding to regenerative cues. MSCs also produce extracellular vesicles (EVs) that carry a range of cargoes, some of which are implicated in cell signalling. Here, we investigate if the changing metabolic state of MSCs alters the cargoes they package into EVs, and how these changing cargoes act on dormant breast cancer cells (BCCs) using an in vitro BCC spheroid model and a scratch assay to create a regenerative demand on MSCs. Our findings show that EVs produced by standard MSCs contain glycolytic metabolites that maintain BCC dormancy. When MSCs are placed under a regenerative demand and increase their respiration to fuel differentiation, these metabolites disappear from the EV cargo and their absence encourages rapid growth in the BCC spheroids. This work implicates EVs in cancer cell dormancy in the bone marrow niche and indicates that pressures on the niche, such as regeneration, can be a driver of BCC activation.

## Introduction

Despite advances in diagnostics, therapies and improvements in patient outcomes, ∼30% of early stage breast cancer (BC) patients have metastatic cancer, which is responsible for most BC deaths (1, 2). Breast cancer cells (BCCs) preferentially metastasise to the bone marrow niche, where they interact with resident mesenchymal stromal cells (MSCs) and can enter dormancy, surviving for decades in the endosteal (bone-lining), region of the marrow (3). Cancer dormancy is defined as a stage during which cells stop dividing and survive in a quiescent state until environmental cues are believed to induce them to proliferate again (4–7). While in quiescence, cells arrest in the G_0_-G_1_ cell cycle phase, which allows metastasised BCCs to avoid detection and treatment, thereby further reinforcing their clinical burden (7). While research into BC dormancy within the marrow has been extensive, little is known about the mechanisms of BCC reactivation.

BCC dormancy in the BM occurs due to the supporting BM niche microenvironment. BCCs enter the BM at the perivascular region, utilising the C-X-C motif chemokine ligand 12 (CXCL12) and C-X-C motif chemokine receptor 4 (CXCR4) signalling axis (8, 9). Once in the BM niche, BCCs communicate with resident MSCs, benefit from the immunosuppressive environment they create, and transition to dormancy. BCC dormancy onset is thought to involve the production of cytokines from perivascular niche MSCs and their gap junctional intercellular communication (GJIC) with BCCs (10, 11). While cytokines, such as CXCL12 and transforming growth factor beta 1 and 2 (TGFβ1 and 2), are implicated in BCC dormancy, there is emerging evidence that MSC-derived extracellular vesicles (MSC-EVs) are also involved in BCC dormancy onset. MSC-EVs carry a variety of cargoes, including the miRNAs miR-126, miR-222 and miR-342-3p, the transfer of which to BCCs is believed to inhibit breast cancer cell proliferation and/or metastasis and to induce dormancy (12–15). MSC-EVs also reportedly have a dose-dependent effect on BCC migration, proliferation and adhesion in 2D and 3D spheroids in vitro, potentially via the Wnt/ β-catenin signalling pathway, initiating a more epithelial phenotype and a net loss of tumourigenicity (16–19).

Depending on the cellular microenvironment, MSCs can exhibit different inflammatory and regenerative behaviours. For example, changes in MSC phenotype are observed in BC patients in an age-dependent manner, after an injury or after intense drug treatment, such as chemotherapy (20–22). In this study, we have developed and characterised a simple MSC wound healing model in vitro to investigate the role of the MSC regenerative phenotype on MSC-EV and their effect on BCC communication and on BCC dormancy and reactivation within BCC 3D spheroids. As noted above, the protein (cytokine, chemokine and growth factor) and nucleic acid (miRNA) cargos of EVs are a current focus of research because of their potential role in BCC dormancy EVs also contain metabolites, and the role of this EV cargo type in BCC dormancy is poorly understood (23, 24), even though bioactive metabolites are known to potentiate cell signalling.

Respiring MSCs undergo oxidative phosphorylation (OXPHOS) as they differentiate and also when involved in regenerating and repairing bone, cartilage and reticular tissues (25, 26). It is possible that self-renewing and quiescent MSCs control this regenerative phenotype via increased oxidative glycolysis (and supressed OXPHOS), in a manner similar to that of the Warburg effect (27), in which cancer cells growing in a glucose-rich environment maintain carbon, lost as CO_2_ during OXPHOS, to build new cells (28, 29). Increased oxidative glycolysis for self-renewal and pluripotency, by suppression of oxidative phosphorylation, has also been observed in pluripotent stem cells (30, 31) and has been linked to the maintenance of the immunomodulatory MSC phenotype (27).

Here, we investigate our hypothesis that exposing MSCs to the demands of wound-healing will shift their respiration from glycolysis to OXPHOS and change the metabolites they package into EVs and that these subsequently altered EV cargoes will either promote BCC dormancy or activation. Specifically, we compared the interactions of a 3D breast cancer cell (MCF7) spheroid model with MSC-EVs isolated from: (a) a standard MSC monolayer, which we propose induces a quiescent BCC phenotype (pQ-EVs); and (b) a regenerative scratch wound MSC model, which we propose activates BCCs (pA-EVs). The reactivation of dormant BCCs at metastatic sites is hypothesised to be a mechanism of both early and late BC recurrence, where the elapsed time after cancer recurrence upon successful treatment depends on the BC subtype (32–35). For instance, triple negative BC patients typically experience cancer recurrence after 1-2 years. By contrast, luminal BC (which presents with estrogen receptor (ER) ^+^ve status) is associated with a much later recurrence, typically ranging from 5-20 years after remission, indicating that ER^+^ve BCCs undergo a lengthy period of dormancy (34, 36). This is supported by the fact that ER^+^ve BC recurrence occurs almost entirely at the site of metastasis (37). The ER^+^ve BC cell line MCF7 was therefore chosen as a representative BCC model for studying BC dormancy/ recurrence.

Our novel findings show that in our 3D model, pQ-EVs promote glycolysis and a dormant BCC phenotype, while pA-EVs confer an active BCC phenotype, relying more on OXPHOS. These changes highlight the influence that the MSC microenvironment has on the MSC phenotype and metabolic statues, and how these changes influence the metabolome cargo packaged into secreted MSCs EVs to bring about changes in BCC activity in vitro that might shed light on how BCCs are reactivated in vitro.

## Results

### MSC-EVs Isolated and Characterised under Regenerative Demand

We placed MSCs into a regenerative environment using a scratch assay, in which a scratch was created across a confluent monolayer of MSCs to mimic a wound; the cells repaired and closed the wound within 44 hours (**supplementary figure 1a, b**). While we observed no change in MSC viability, the MSC differentiation marker CD105 showed significantly increased expression in scratched MSCs during the scratch-recovery period (**supplementary figure 1c-e; supplementary table 1**). CD105 is linked to transforming growth factor beta (TGF-β)-mediated regulation of MSC differentiation and lineage commitment (38). We also observed increased expression of migration markers in the scratched cells, including of matrix metalloproteinase 2 (MMP2), RhoA, Rac 1, collagen type 1 alpha 1 chain (Col1A1) and TGF-β (39–41), peaking at 6 hours post scratch, with a parallel increase in the cytokine secretome over the 48 hour scratch-recovery period (**supplementary figure 1f, g**). These findings show that the regenerative demand created in this assay generates a distinct change in MSC phenotype.

**Figure 1.**
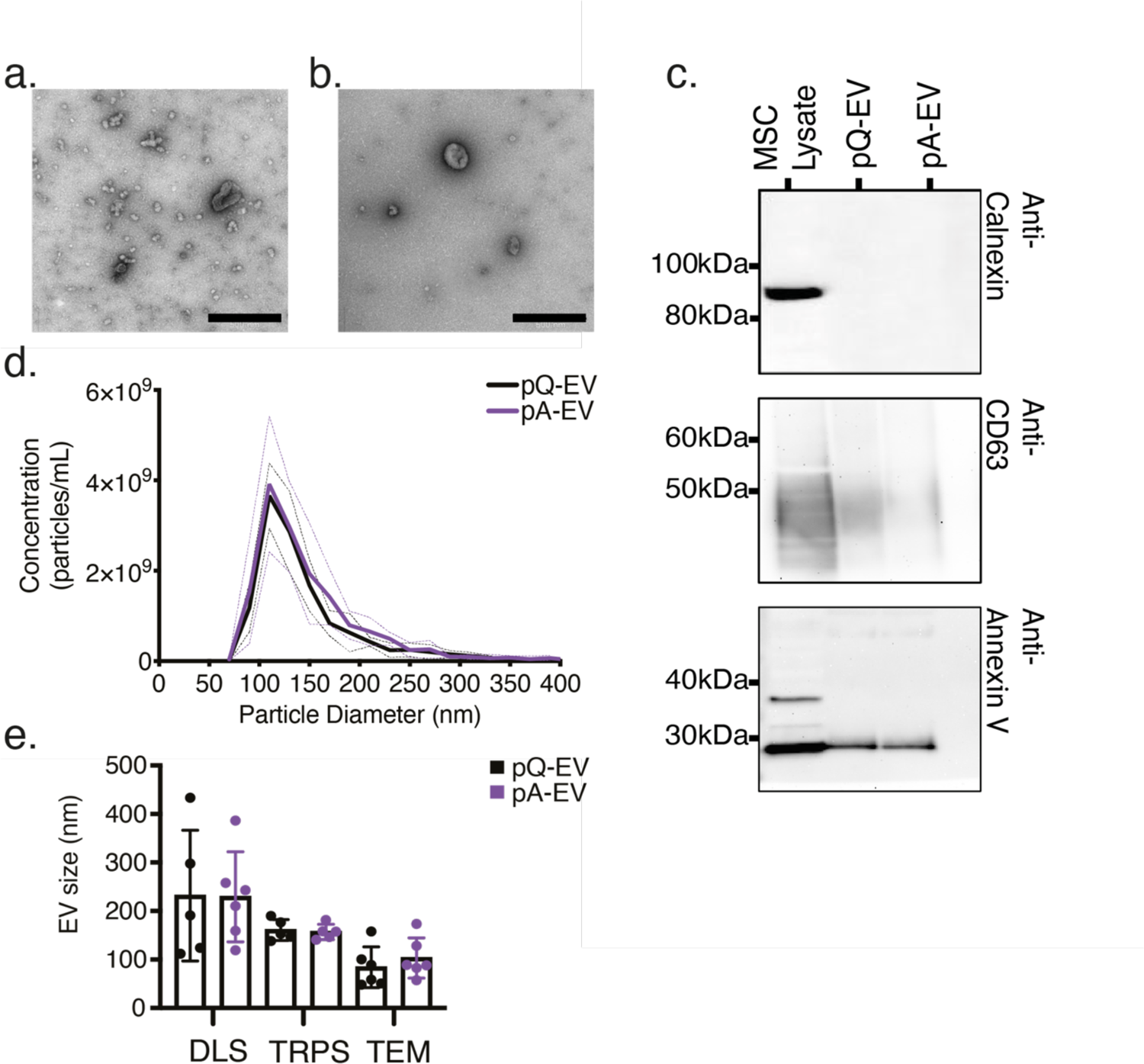
MSC-EV characterisation. MSC-EVs were isolated from pQ-MSC and pA-MSC cultures via differential ultracentrifugation and defined using the MISEV EV characterisation criteria (47). **a, b.)** Transmission electron microscopy (TEM) images confirm the presence of spherical vesicles in isolated samples (scale bar 500 nm). **a**.) pQ-EV, **b**.) pA-EV. **c.)** Western blotting confirmed the presence of positive EV markers CD63 and annexin V, and the absence of negative marker, calnexin. **d.)** Tuneable resistive pulse sensing (TRPS) technology (qNano IZON system, USA) was used to determine the size distribution and concentration of particles in the MSC-EV isolations. Dotted lines represent ± SD. **e.)** A comparison of mean EV size ± SD, quantified via dynamic light scattering (DLS), TRPS and TEM, n=6 isolations, TRPS data from n=3.

**Table 1.**
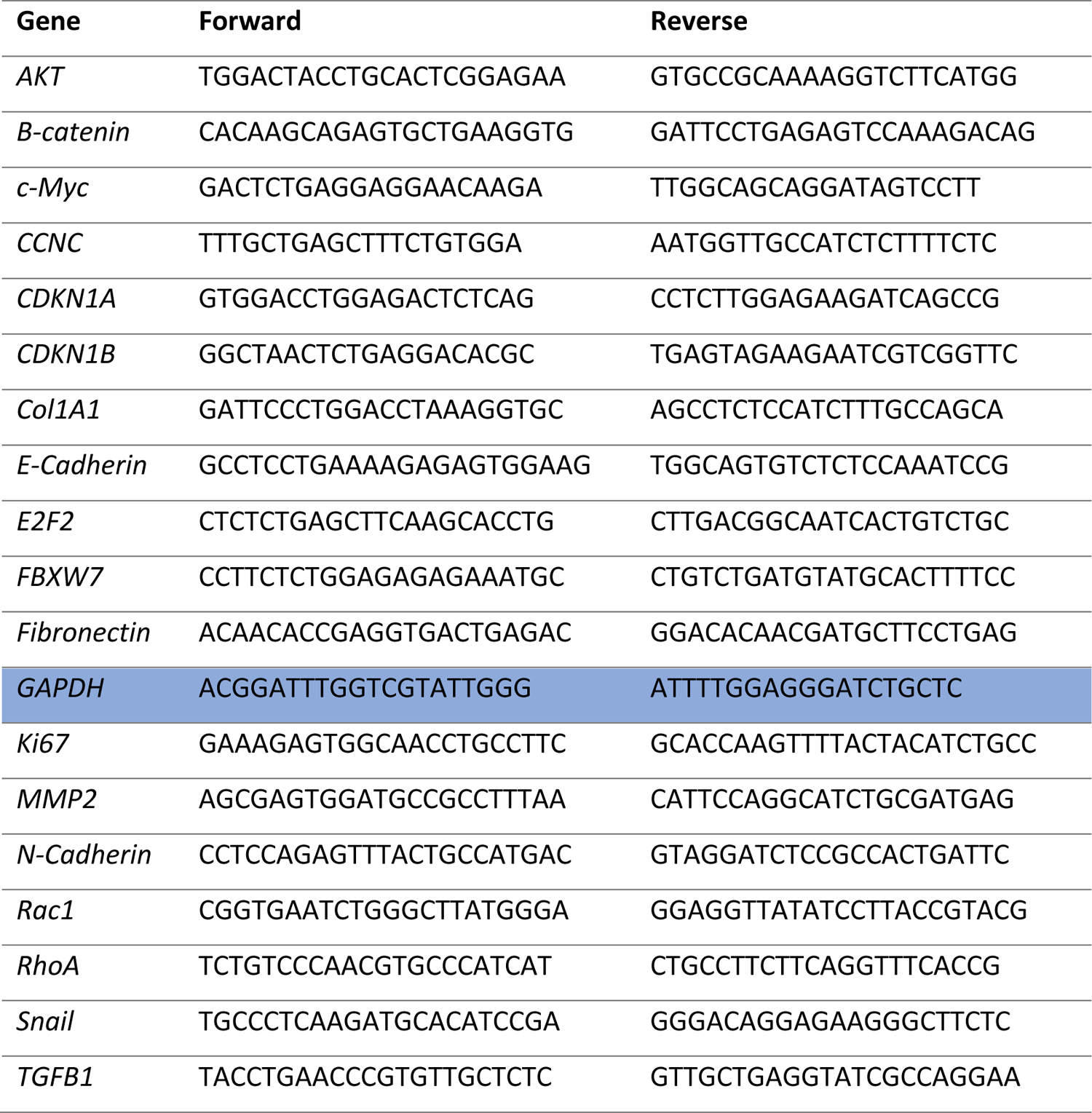
RT-qPCR primer sequences used in RT-qPCR studies Shaded cells indicate housekeeping control gene used for normalisation.

Next, we isolated MSC-EVs from both unscratched and scratched cultures via differential ultracentrifugation and characterised them (42–47). We observed no significant differences in EV morphology, size and concentration between the proposed-quiescent (pQ-EV, taken from unscratched cultures) and proposed-activating (pA-EVs, taken from scratched cultures) EV populations (**figure 1**). A relatively pure isolation was obtained for both populations, with EVs predominantly in the range of 70-150nm (exosome and microvesicle range). In accordance with the updated 2018 minimal Information for Studies of Extracellular Vesicles (MISEV2018) guidelines (47), we refer to these structures as EVs (47, 48).

### MSC-EV Cargo is significantly altered under MSC regenerative demand

To investigate whether MSC-EVs cargoes are altered upon scratch injury, we performed two EV cargo screens. A mass-spectrometry-based untargeted metabolomic screen identified that the metabolic relative abundance is higher in pQ-EVs relative to pA-EVs, indicating that pQ-MSCs package more metabolite cargo into EVs than do pA-MSCs (**figure 2a,b**). We identified the metabolites that showed the greatest differences between the two EV populations. We found that the top six metabolites depleted in pA-EVs relative to pQ-EVs were adenine, adenosine, isonicotinic acid, pyruvate, (R)-lactate and D-glucose, which are all involved in carbohydrate metabolism (specifically glucose pathway signaling) (**figure 2c**). To identify similarities/ differences in metabolites at the pathway level, the top 30 metabolite list was run through Qiagen’s Ingenuity Pathway Analysis (IPA) software. We used the predictive modelling function, where metabolite patterns are linked to biochemical pathways to predict up- or down-regulation of the linked pathway. Fold-change values of pA-EV vs pQ-EV peak intensities were run, subject to a comparison analysis, and the top network pathway was mapped (**figure 2d**). This software uses the literature to derive what it thinks the parent cells would be doing given a particular pattern of metabolite depletions. In our analysis, it predicted the activation of the extracellular signal related kinase (ERK 1/2) and growth hormone pathways, which are both centrally involved in the upregulation of cell growth and proliferation (49, 50). We therefore conclude that in the scratched-assay MSCs, metabolites are being used to drive regenerative responses and so are not available for EV packaging.

**Figure 2.**
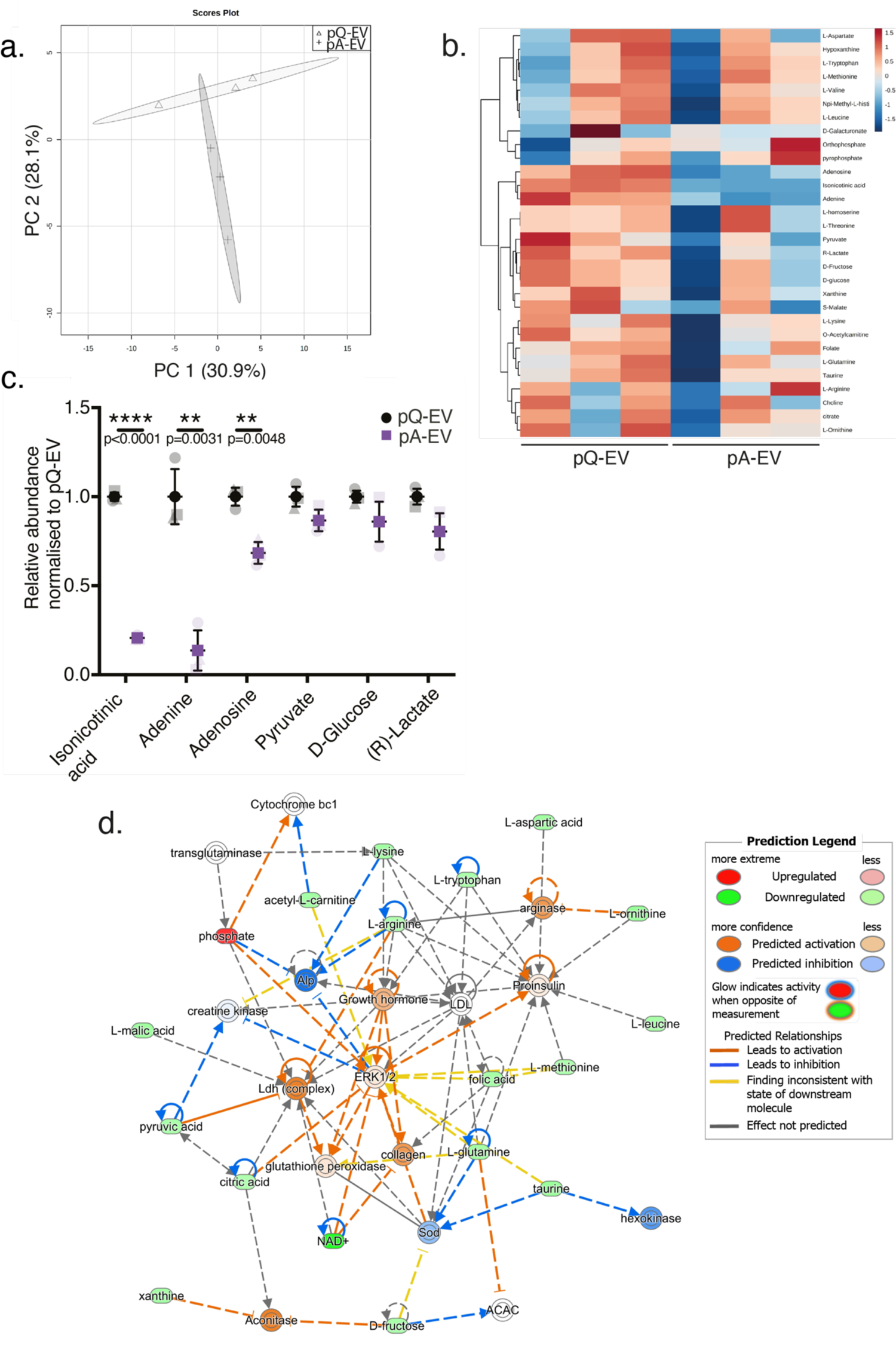
Metabolic analysis of MSC-EV cargo reveals differing metabolic EV signatures dependent on MSC state. The metabolic cargos of pQ-EV and pA-EV samples were investigated using liquid chromatography-mass spectrometry analysis. **a.)** The abundance of each metabolite detected (peak intensities) was subject to principle component analysis (PCA). Each point represents 1 replicate and ellipses represent the spatial borders associated with each condition with a 95% confidence interval, n=3. **b**.) A heatmap of the 30 identified metabolites shows their relative abundance in pQ-EV vs pA-EVs, clustered through average linkage, showing differences between pQ-EV and pA-EV cargo. Metabolites were categorised as either identified or annotated with fragment data, and were filtered using the PIMP software designed by Glasgow Polyomics. Any metabolites showing a fold-change >1.2, in addition to PIMP identified labelling, were further analysed. **c**.) Metabolites that show the greatest differences in peak intensities between pQ-EV vs. pA-EV. These metabolites are involved in carbohydrate metabolism. Mean intensity was normalised to pQ-EV ± SD. An unpaired t-test was performed to assess the statistical difference between samples; ***= p< 0.0001; **= p< 0.01. **d**.) Fold-change values of pA-EV vs pQ-EV peak intensities were analysed using the Comparison Analysis in Ingenuity pathway analysis (IPA, Qiagen) and the top network pathway mapped.

To investigate the proteins packaged into each MSC-EV population, we conducted a mass spectrometry-based and label-free proteomic analyses (**figure 3a**). We then used PCA to analyse the abundance of every protein per sample and its peak intensity. PC1 represents 39.5% and PC2 26% of the observed variance between conditions. The projection of substrate conditions onto PC1 and PC2 separates their proteomic signatures into clear, distinct clusters, indicating that the two MSC populations (pQ-EV and pA-EV) produce discrete proteomic cargo (**figure 3b**). This proteomic EV screen identified a total of 210 hits, of which 27 presented a significant difference between the pQ-EV and pA-EV populations. A pQ-EV vs pA-EV fold-change analysis demonstrated that most of the significant hits were upregulated in pQ-EVs. A heatmap of the top 25 significant hits showed the relative abundance of proteins between pQ-EV and pA-EV, clustered through average linkage (**figure 3c**). With the exception of ecto-ADP-ribosyltransferase and mitogen-activated protein kinase 12-binding inhibitory protein 1 (MBIP1), the significant hits were more abundant in the cargo of pQ-EV MSCs, indicating a higher relative protein abundance in pQ-EV MSC cargo, relative to pA-EV cargo.

**Figure 3.**
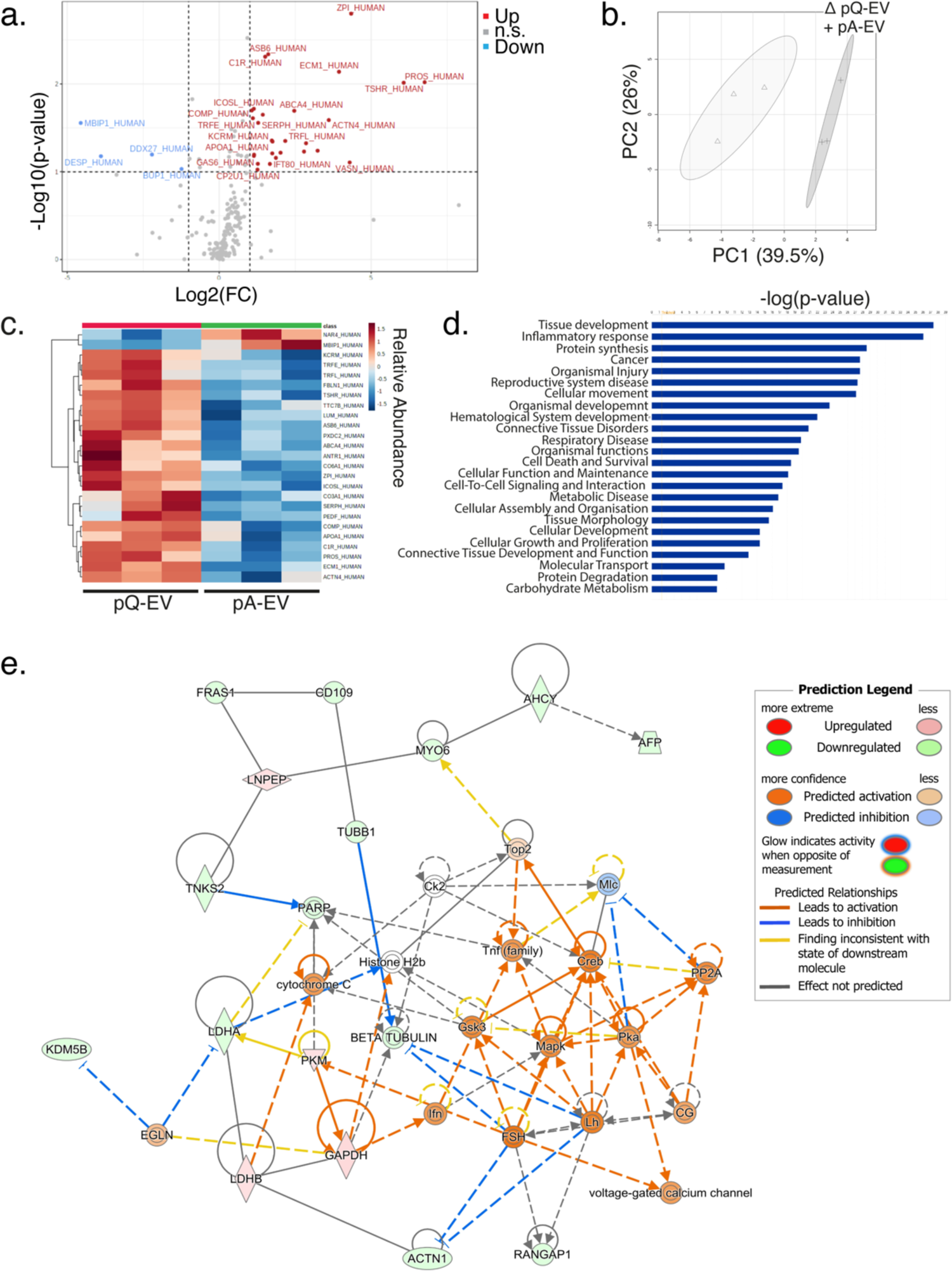
Proteomics screen reveals differences in MSC-EV cargo, with pA-EV cargo linked to cancer cell growth, migration, invasion and metabolic pathways The proteomic cargos of pQ-EV and pA-EV samples were investigated using liquid chromatography-coupled tandem mass spectrometry (LC-MS/MS) analysis, n=3. A.) Fold-change of pQ-EV vs pA-EVs was Log2 transformed and mapped against -Log10(p-value) to show significantly up- and down-regulated proteins identified in the screen. B.) PCA in which each point represents 1 replicate and ellipses represent the spatial borders associated with each condition with a 95% confidence interval. C.) A heatmap showing the top 25 differentially abundant proteins identified between pQ-EV and pA-EVs, clustered through average linkage. D.) pA-EV vs pQ-EV normalised abundance fold-change values comparatively analysed using Ingenuity pathway analysis (IPA, Qiagen). The main biological processes affected are shown. e.) Normalised peak intensities fold-change values of pA-EV vs pQ-EV were also analysed using (IPA) and the top 2 network pathways mapped, n=3. E.) Mitogen-activated protein kinase (MAPK) and tumour necrosis factor (TNF) family pathways were predicted to be activated.

We also conducted a comparison analysis of the pA-EV vs pQ-EV normalised abundance fold-change values using IPA to investigate the main biological processes affected (**figure 3d**). As before, the software analyses the protein abundance data in light of what the literature indicates parent cells would be doing given this pattern of protein depletion. This analysis predicted numerous wound healing processes to be more affected in pA-EVs, namely; tissue development, inflammatory response, protein synthesis, organismal injury, cellular function and maintenance, cellular assembly and organisation, cellular development, cellular growth and proliferation, connective tissue development and function. Given these results, we propose that unwounded MSCs in our assay are not depleting these proteins as they have no demand for them; the proteins are thus packaged into EVs. Previous MSC-EV proteomic studies have also linked the MSC-EV cargo to the parent cell (51, 52). The protein abundance pattern in pQ-EV MSCs relative to pA-EV also indicates that if this depletion pattern was seen in the parent cells, that carbohydrate metabolism was being influenced, indicating an increased energy demand in the scratched-assay MSCs.

We also performed a comparison analysis on the MSC-EV proteomic data and mapped the top two network pathways (**figure 3e and supplementary figure 2**). Network 1 depicts that the mitogen-activated protein kinase (MAPK) and tumour necrosis factor (TNF) family pathways are predicted to be activated in the parent cells based on the protein cargo depletion patterns in pA-EV vs pQ-EV (**figure 3e**). This data supported the IPA analysis of metabolomic depletions, which identified and predicted an activation of the MAPK/ ERK 1/2 pathway. Furthermore, cAMP (cyclic adenosine monophosphate), protein kinase A (PKA) and cAMP-responsive element binding protein (CREB) are predicted to be up-regulated in the pA-EV parent cells based on these depletion patterns; all of these pathways regulate cancer cell growth, migration, invasion and metabolism (53).

Evidence suggests that ERK 1/2 activation can determine whether cancer cells proliferate or enter a state of dormancy (61). Cancer cells in a constantly proliferating state exhibit constitutive ERK 1/2 activation, permitting G_0_-G_1_ to S phase transition and leading to cell division (62, 63). Notably, low ERK 1/2 activation is recognised as a general mechanism that underlies cancer cell dormancy (61). P38/ ERK 1/2 activity thereby provide a signalling balance that regulates cancer cell fate.

Network 2 (**supplementary figure 2**) depicts growth and inflammatory response pathways. Here, IPA predicts that the depletion pattern from pQ-EVs to pA-EVs, if in a parent cell, would lead to the activation of vascular endothelial growth factors (VEGF), Interferon-Beta (INF-β), RNA polymerase II and nuclear factor kappa B (NFκB) family networks. As the pA-MSCs parent cells are under regenerative demand, an increase in these pathways is logical, with increases in protein translation and inflammatory response commonly seen during wound healing (54–56).

Under cellular stress conditions, metabolic demand for nutrients and for cellular building blocks increases, which may partly explain the packaging trends seen in our screens, in which pA-MSCs need to retain metabolites and proteins for their own repair (57). This is likely linked to our observed depletion of metabolites and proteins involved in cell growth and wound healing in EVs derived from activated MSCs. We know that in the endosteal and perivascular BM niches that MSCs self-renew and retain stem cell phenotype (30, 31), and that BCCs enter dormancy there. This allows us to hypothesise that the more abundant cargos of the pQ-EVs should not activate the BCCs and may even potentiate dormancy.

### pQ-EVs differentially influence 3D MCF7 spheroid activity compared to pA-EVs

We used MCF7 cells to investigate the effect of MSC-EVs on breast cancer cells by developing a 3D MCF7 spheroid culture model. We began by confirming the viability of this model and MSC-EV uptake by spheroids (**supplementary figure 3**). We then investigated the effects of MSC-EVs on MCF7 spheroid proliferation, migration and EMT. MCF7s were initially seeded at a very low density, mimicking the number of cells that would arrive in the BM upon BC metastasis (58). We then performed a colony formation assay (CFA) (**figure 4a,b**) for 2 weeks, supplementing each culture with either pQ-EVs or pA-EVs every three days. pQ-EV treated-colonies formed statistically significantly fewer colonies relative to both untreated (UT) and pA-EV-treated colonies (**figure 4a, b**). No significant differences in colony formation were observed between UT and pA-EV treated samples (**figure 4a, b**). We next performed RT-qPCR to assess the expression of several dormancy- and activity-related genes following EV treatment (**figure 4c**). Dormancy-related genes showed increased expression 24 hours after spheroids were treated with either pQ-EVs or pA-EVs, 80% of which had a higher expression upon treatment with pQ-EVs vs pA-EVs. Two cancer cell proliferation genes, *c-MYC* and *Ki67*, were downregulated 24 hours after treatment with pQ-EV, but were increased in pA-EV treated spheroids at 24 hours. In treated spheroids, a global downregulation of all genes was observed 72 hours after treatment compared to UT spheroids. Ki67 is an established proliferation marker that is often used as a prognostic marker for BC (59, 60). Ki67 is expressed in the nuclei of cells in all phases of the cell cycle, except for dormant cells resting in the G_0_ phase (61, 62). This characteristic expression thus makes Ki67 an excellent marker for determining the proliferative activity of a culture. In pQ-EV treated MCF7 spheroids, Ki67 levels were decreased at both 24 and 72 hours post treatment, as assayed by Western blot, relative to both UT and pA-EV treated cultures at these timepoints (**figure 4d,e**). pA-EV treated spheroids expressed significantly higher Ki67 levels compared to pQ-EV treated spheroids. This data suggests that pQ-EV treatment decreased the proliferation rate of spheroids whilst pA-EV treatment increased their proliferation. We also used propidium iodide staining to assess whether these changes in proliferation were reflected in altered DNA content in MCF7 spheroids (**figure 4f,g**). We observed a significant increase in the proportion of cells in G_1_ and G_2_/M 24 hours after pQ-EV treatment relative to UT samples, and a significant increase in the proportion of cells in G_1_ 24 hours after pA-EV treatment relative to UT samples (**figure 4f**). In pQ-EV treated spheroids, we observed a significant increase in cells in G_1_ and a decrease in cells in S phase 72 hours after treatment, compared to both pA-EV treated and UT spheroid cultures at this timepoint (**figure 4g**). These findings indicate that pQ-EV treatment caused a marked reduction in MCF7 spheroid proliferation at the 72 hour timepoint compared to treatment with pA-EVs. 72 hours of pQ-EV treatment also caused a significant increase in the number of MCF7 spheroid cells in G_1_, but not in G_2_/M. This suggests that pQ-EV treated cells, which had divided by 24 hours, did not re-enter the cell cycle and remained in cell cycle arrest.

**Figure 4.**
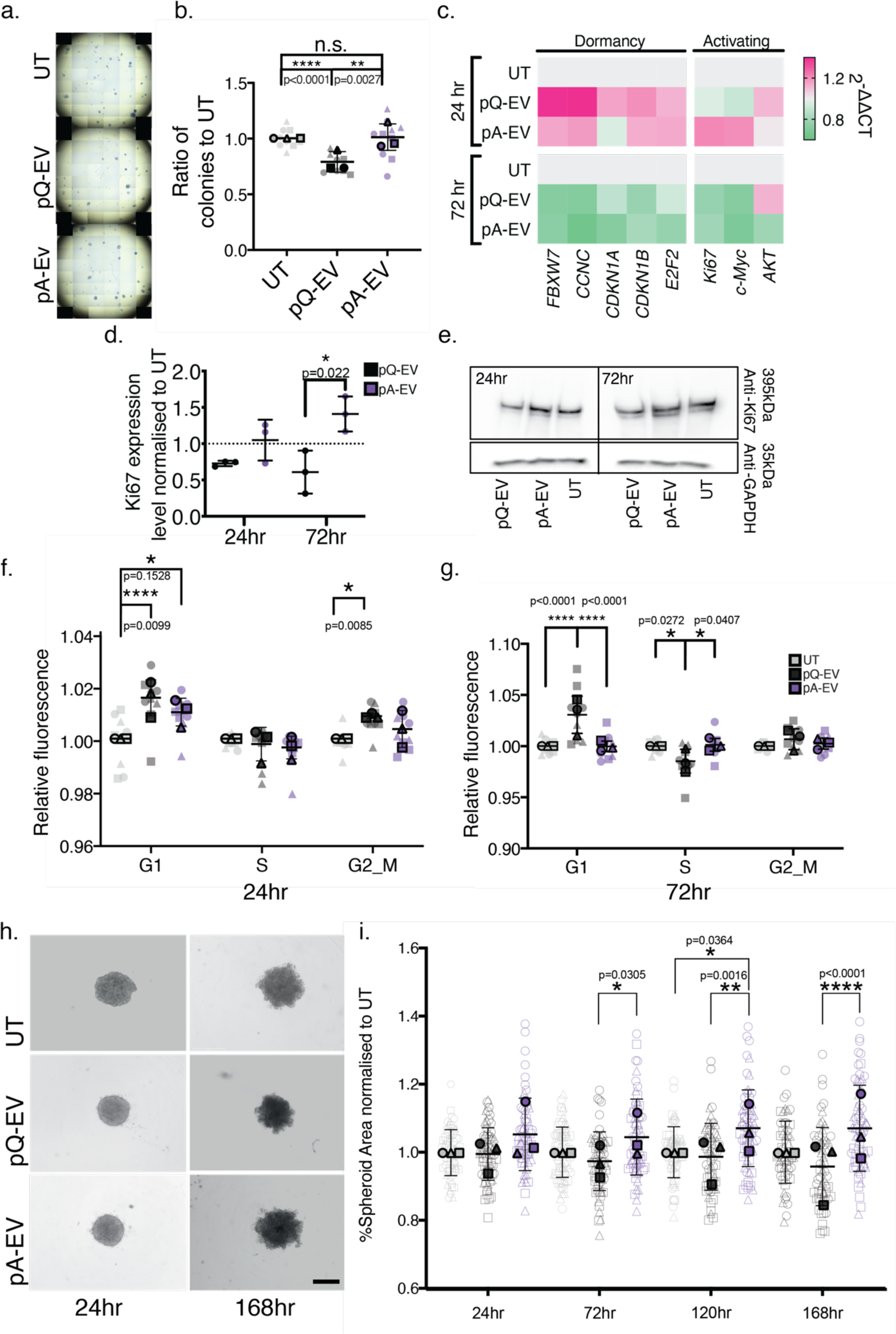
MSC-EVs differentially influence 3D MCF7 spheroid activity. **a, b**.) Colony formation assay used to assess the effect of MSC-EV treatment over 2 weeks. MCF7s were seeded at a low density in 2D monolayer. Cultures were either untreated (UT) or treated with pQ-EV or pA-EV every 3 days for 2 weeks, after which colonies were fixed, stained and counted. MSC-EV treatments were normalised to the UT control. **a**.) Representative wells with stained colonies. **b**.) Mean colonies per condition ± SD plotted, and an unpaired t-test performed, **p<0.01, ****p< 0.0001, n=3. **c**.) MCF7 spheroid cultures were untreated (UT) or incubated for 24/ 72 hours with pQ-EV or pA-EV before RNA extraction for cell cycle gene expression analysis. Expression data for each gene was normalised to housekeeping controls, then normalised to UT cells to generate 2^-ΔΔCT^ values. Selected genes expressed during dormant and active cell cycle phases were investigated: upregulated genes (pink), downregulated genes (green). 3 technical repeats performed per independent experiment, n=3. **d**.) Ki67 band density was normalised to a GAPDH loading control and then further normalised to UT samples to calculate a fold-change value. Graph shows mean ± SD and statistical analysis was run on the mean fold change value calculated from 3 independent experiments using an unpaired t-test, n=3. **e.**) Representative Western blot (one of 3) showing Ki67 levels in MCF7 spheroid cultures incubated for 24/ 72 hours with pQ-EV/ pA-EV relative to UT controls. **f.**) 3D MCF7 spheroids treated with pQ-EV/ pA-EV or UT and incubated for **f**.) 24 or **g**.) 72 hours. Spheroids were then collected, dissociated with collagenase D, washed, fixed and stained with FxCycle™ PI/RNase Staining Solution. The stained single cell suspensions then underwent flow cytometry. Graphs represent geometric mean fluorescent intensity per cell cycle phase, normalised to UT cells ± SD, with a minimum of 3000 cells analysed per 3 technical repeats. A 2-way ANOVA with multiple comparisons test was done, *= p<0.05, ****= p<0.0001, n=3. **h-i.**) Effect of MSC-EV treatment on MCF7 spheroid size, migratory projections and overall proliferation. At 24, 72, 120 and 168 hours of UT/ pQ-EV/pA-EV treatment, spheroids were imaged using the EVOS M7000 Imaging system. A minimum of 20 spheroids were imaged per treatment and the area of each spheroid measured via ImageJ. **h**.) Representative images of pQ-EV-, pA-EV treated and UT spheroids at 24 and 168 hour timepoints. Scale bar represents 650 µm. **i**.) Mean spheroid area ± SD plotted and a one-way ANOVA followed by a Kruskal Wallis multiple comparisons test was done, *= p < 0.05, **= p< 0.01, ****= p< 0.0001, n=3.

To investigate the influence of MSC-EV treatments on MCF7 migration, we performed: (i) a 2D migration assay via the Ibidi µ-slide chemotaxis protocol, which enables the directional motility of cells to be observed and measured in response to a specific chemoattractant; and (ii) a 3D spheroid growth assay, to analyse spheroid migratory projections and the overall proliferative effect of these treatments (**supplementary figure 4** and **figure 4h,i**). Over a 48-hour treatment period, the accumulated distance and velocity of pA-EV treated MCF7 cells was higher than that of UT cells and significantly higher than that of pQ-EV treated cells. These findings indicate that pQ-EV treatment results in decreased MCF7 motility, while pA-EV treatment produces a more chemoattractant and migratory MCF7 signal (**supplementary figure 4**). At 24, 72, 120 and 168 hours of culture with pA-EV or pQ-EV, spheroids were imaged and their area measured. Significant differences were noted after 72 hours of EV treatment, with pQ-EV-treated spheroids presenting a smaller area compared to pA-EV-treated spheroids (**figure 4i**). Representative images show that at 24 hours, spheroids do not exhibit many projections, but that projections become apparent after 72 hours, especially in UT and pA-EV-treated samples (**figure 4h**). We used RT-qPCR to investigate the expression of a selection of 3D MCF7 EMT markers following the MSC-EV treatment (**supplementary figure 5**). Very few significant differences in these markers’ expression were observed; the main observation being that pQ-EV treatment caused a significant downregulation in the mesenchymal marker *SNAIL* at 72 hours of treatment compared to UT spheroids. This finding suggests that this treatment might cause a slight loss in mesenchymal phenotype, as would be expected with less migratory cancer cells (63).

**Figure 5.**
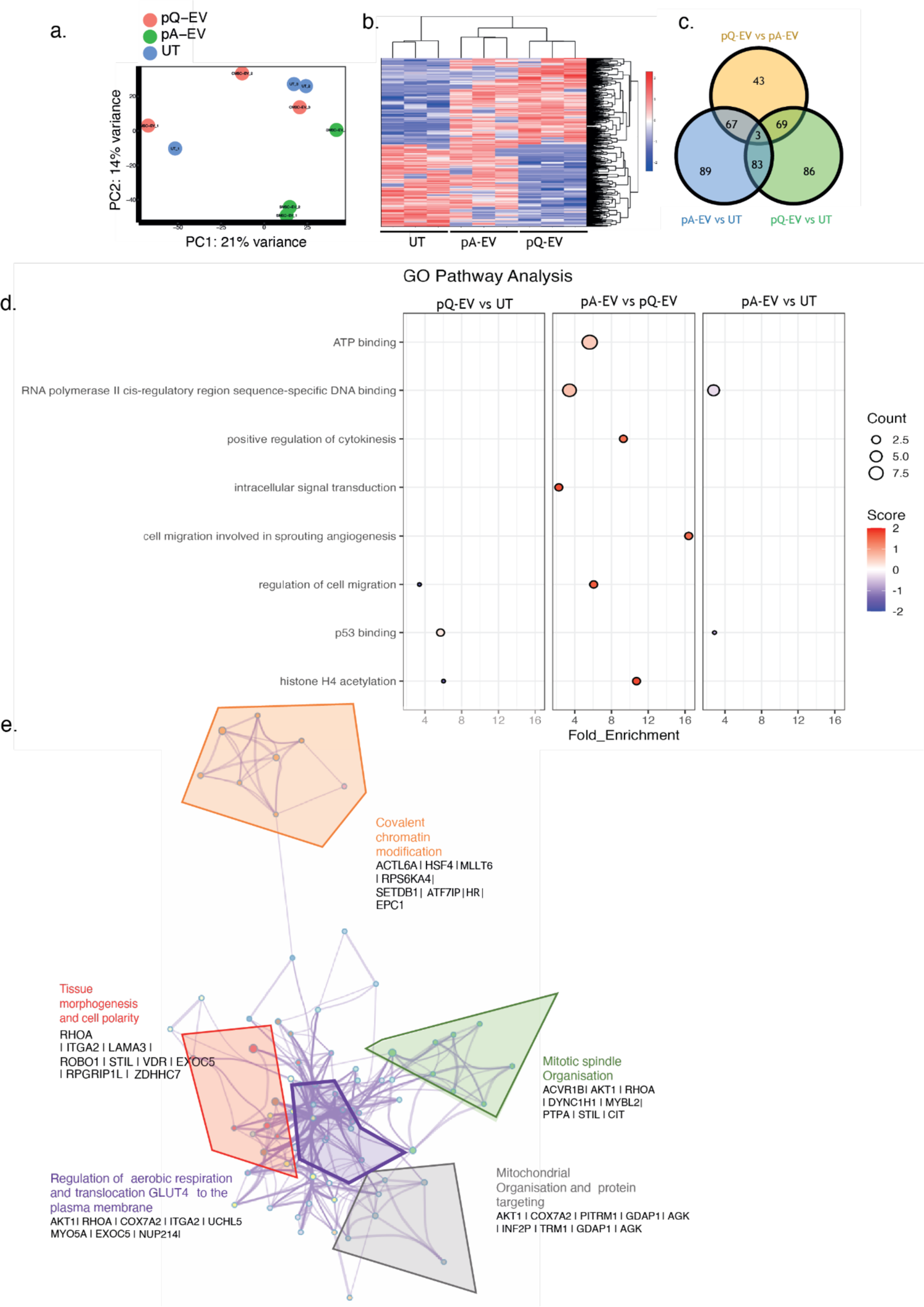
MCF7 spheroid cultures show significantly different RNA-seq profiles 24 hours after treatment with pQ-EV or pA-EV. MCF7 spheroid cultures were untreated (UT) or incubated for 24 hours with pQ-EV or pA-EV before RNA was extracted for NGS RNA sequencing. **a**.) Principle component analysis in which each point represents 1 replicate, n=3. **b**.) Heatmap showing all hits, clustered through average linkage. **c**.) Venn diagram showing the distribution of all significant hits (p<0.001), across the different conditions. **d**.) Selected hits from GO pathway enrichment. **e.**) Network diagrams illustrate pathways affected, where each node represents a significantly enriched term. Node size is proportional to the number of contributing genes. Similar terms with a high degree of redundancy were clustered, as depicted, including a description and the genes involved. Map shows that pA-EV-treated spheroid pathways are significantly upregulated compared to pQ-EV-treated spheroids.

### MCF7 spheroid RNA sequence profiles change with MSC-EV treatment

To investigate if MSC-EV treatment altered the transcriptional profile of treated MCF7 spheroids, we performed RNA-seq at 24 and 72 hours post treatment (volcano plots **supplementary figure 6**). The PC, heatmap and Venn diagram figures in **figure 5a-c** show a distinct MCF7 spheroid RNA seq profile for each treatment condition at the 24 hour timepoint. At 24 hours after pA-EV treatment, the PCA plot shows that MCF7 cells have a distinct profile relative to UT or pQ-EV treated MCF7 spheroids (**figure 5a**). UT and pQ-EV treated MCF7 spheroid cells also have different transcriptional profiles, as shown by heatmap analysis, and so are distinct from each other. MCF7 spheroids treated with pA-EVs exhibit a transcriptional profile that is indicative of general up-regulation (**figure 5b**). Venn diagram analysis shows that more changes in this gene set were identified in the MSC-EV treated MCF7 spheroids, irrespective of whether they were treated with pQ-EVs or pA-EVs (**figure 5c**). Several key findings emerged from our GO pathway analysis of the 24 hour RNA seq data, for example, energy pathways (ATP binding), transcription, cell division and migration pathways are more highly up-regulated in pA-EV-than in pQ-EV-treated MCF7 spheroids (**figure 5d**). Very few differences were noted for either pQ-EV- or pA-EV-treated MCF7 spheroids relative to UT spheroids, likely because fewer transcript sets are differentially regulated in the UT MCF7 spheroid population (**figure 5d**). This pattern of pA-EVs producing a more active transcriptional pattern in MCF7 spheroids was also noted after 72 hours of culture (**supplementary figure 7**). Network mapping of selected transcriptional changes again highlighted that pA-EV-treated MCF7 spheroids showed increased transcriptional activity relative to pQ-EV-treated and UT spheroids. Noteworthy areas of transcriptional activation in pA-EV-treated MCF7 spheroids included metabolic and cell growth pathways, and specifically chromatin modification, tissue morphogenesis, mitotic spindle organisation, mitochondrial organisation, and regulation of aerobic respiration (**figure 6e**) (64–66). Together, these patterns of expression indicate that pA-EVs elicit the transcription of genes involved in MCF7 spheroid activity.

**Figure 6.**
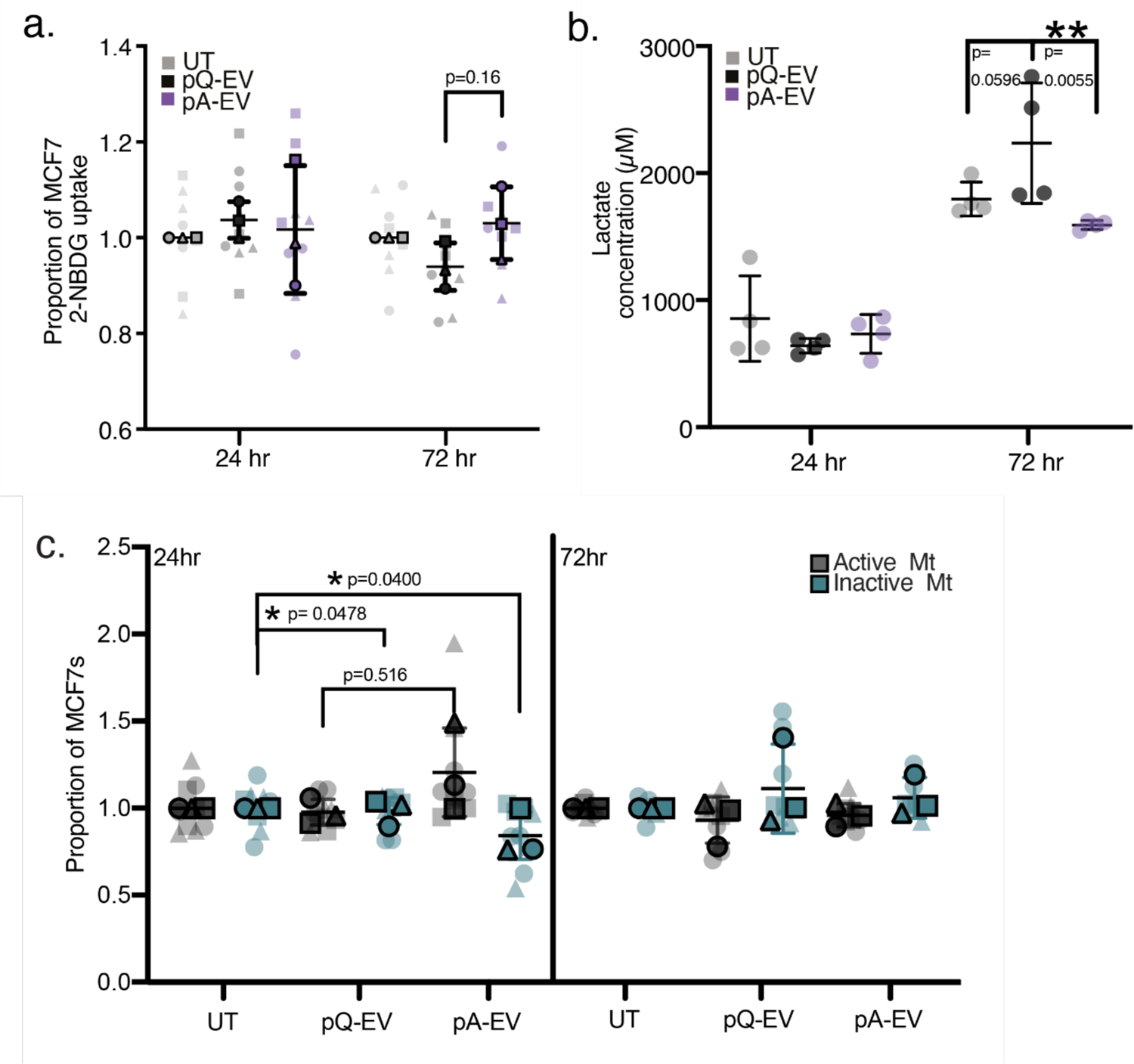
Metabolic assays reveal differing effects of pQ-EV and pA-EV on MCF7 spheroid cultures. We assessed the influence of pQ-EV and pA-EV on MCF7 spheroid culture metabolism using various metabolic assays. MCF7s were UT or incubated for 24 or 72 hours with pQ-EV or pA-EV for each assay. **a**.) Glucose uptake by MCF7 spheroid cultures assessed using the glucose analogue 2-NBDG and flow cytometry. Mean fluorescent intensity was normalised to UT cells. Graph presents the mean proportion of 2-NBDG uptake ± SD, with 3 technical repeats and minimum of 5000 cells measured per sample. A 2-way ANOVA with multiple comparisons was performed, n=3. **b**.) MCF7 spheroid culture lactate secretion levels were measured, to assess cellular levels of glycolytic metabolism. Supernatants were collected at each timepoint, and lactate levels quantified using the Lactate Glo^TM^ Assay (Promega, J5021). Graph depicts mean ± SEM concentration, with 5 technical repeats per sample. A 2-way ANOVA statistical test was performed, *= p<0.05, n=1. **c**.) Mitochondrial respiration was assessed using the JC-1 dye. Active mitochondria accumulate red fluorescence, inactive mitochondria remain green. Quantification of red and green fluorescence by flow cytometry provides a functional readout of mitochondrial activity. Graph depicts mean ± SEM proportion of active and inactive mitochondria for each condition, normalised to UT cells. 3 technical repeats with a minimum of 5000 cells were measured per sample and an unpaired t-test completed between each condition, n=3.

**Figure 7.**
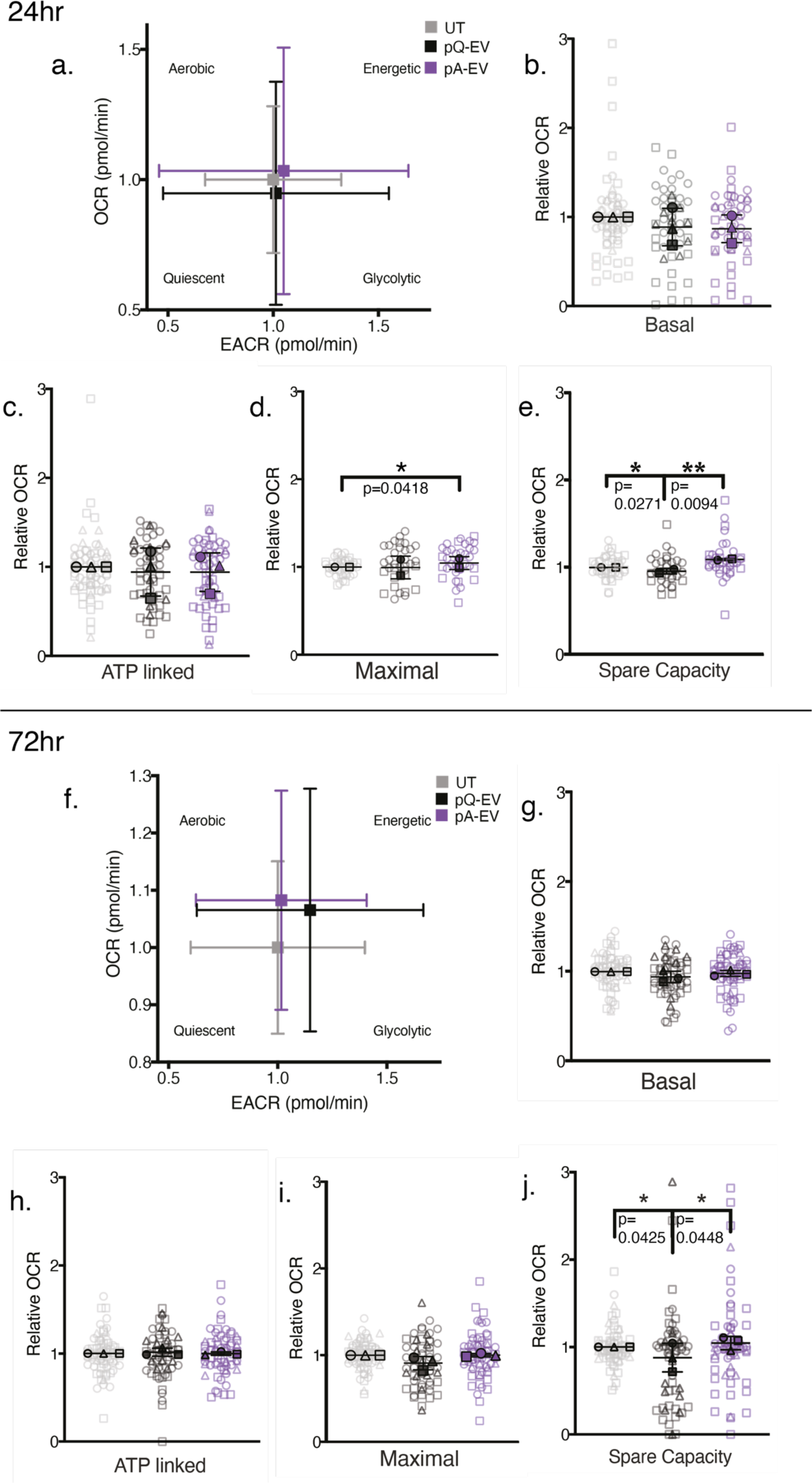
Seahorse respiratory analysis reveals MCF7s in spheroid culture shift to a quiescent phenotype on pQ-EV treatment. The mitochondrial bioenergetics of MCF7 spheroid cultures in response to pQ-EV or pA-EV was assessed using the Seahorse assay. MCF7s were untreated (UT) or incubated for **a-e.**) 24 or **f-j.)**72 hours with pQ-EV or pA-EV. Spheroids were then treated sequentially with oligomycin (complex V inhibitor), FCCP (a protonophore) and antimycin A/ rotenone (complex III inhibitor/ complex I inhibitor), and oxygen consumption rate (OCR) and extracellular acidification rate (ECAR) were measured over time. **a, f.**) Energy maps highlight the overall metabolic phenotype of cultures after **a**.) 24 hours or **f.**) 72 hours of treatment. Graph shows mean ECAR vs OCR ± SD, n=3. **b-e.**) Graphs represent the mean OCR levels ± SD, normalised to UT cells after 24 hours of treatment, including a minimum of 6 spheroids analysed per sample. An unpaired t-test was performed between each sample for all conditions, *= p<0.05, **= p<0.01, n=3. **g-j**.) Graphs represent the mean OCR levels ± SD, normalised to UT cells after 72 hours of treatment, including a minimum of 6 spheroids analysed per sample. An unpaired t-test was performed between each sample for all conditions, *= p<0.05, **= p<0.01, n=3

### MSC-EV treatment differentially influences respiration in MCF7 spheroids

The deregulation of cellular energetics is a hallmark of cancer (64). Yet, research into the metabolic properties of dormant BCCs is limited, and the studies performed to date are contradictory (35). For example, several reports have concluded that a shift in BCCs towards glycolysis is responsible for BCC dormancy, whereas others have reported that a BCC shift towards OXPHOS induces dormancy (35,67– 70). Notably, none of these studies explained a metabolic mechanism for BCC recurrence.

To support their high rates of proliferation, BCCs maintain high glucose uptake (71). We therefore assessed glucose uptake in MCF7 spheroids that were either UT or treated with MSC-EVs (**figure 6a**). Specifically, we treated MCF7 spheroids for 24 and 72 hours, and assessed the uptake of the fluorescent glucose analogue 2-NBDG using flow cytometry analysis. After 24 and 72 hours of treatment, we observed no significant difference in glucose uptake between the UT and the pQ-EV- or pA-EV-treated conditions. However, we observed a significant decrease in the levels of glucose uptake when we compared the pQ-EV-treated to the pA-EV-treated MCF7 cultures at 72 hours. This finding indicates that after 72 hours, pQ-EV-treated cells decrease their energetic demand. Simultaneously, lactate levels increased in the pQ-EV-treated MCF7 spheroids after 72 hours of treatment (**figure 6b**), which is indicative of enhanced aerobic glycolysis within the cytoplasm, as seen in the Warburg effect (in which lactate is produced with glucose consumption) (72). This data supports that pQ-EVs caused an increase in MCF7 spheroid aerobic glycolytic metabolism compared to UT and pA-EV treated cultures. Conversely, pA-EVs demonstrate a trend for increased glucose consumption alongside reduced lactate at 72 hours (figure 7a, b), supporting enhanced OXPHOS within the mitochondria.

Changes in glycolysis / OXPHOS balance should affect mitochondria activity. To investigate this, we assayed for mitochondrial activity using the dye JC-1 and flow cytometry (**figure 7c**). We assessed the cellular red:green JC-1 fluorescent ratio to evaluate the proportion of active vs inactive mitochondria in 3D MCF7 cultures. We observed an increase in the level of active mitochondria and a significant decrease in the level of inactive mitochondria 24 hours after pA-EV treatment. Compared to UT and pQ-EV treatment, pA-EV treatment appeared to upregulate OXPHOS levels in MCF7 cultures, which then return to UT levels after 72 hours.

Together, these data indicate that pQ-EVs produce an oxidative-glycolytic phenotype in treated MCF7 spheroids, supporting the earlier proliferation and transcriptomic data, which shows that these cells are less active. By contrast, pA-EVs confer a more OXPHOS-biased metabolism along with a more proliferative phenotype in MCF7 spheroids.

### pQ-EVs confer a quiescent whilst pA-EVs confer an energetic phenotype

To gain a more detailed insight into the influence of MSC-EVs on MCF7 spheroid mitochondrial bioenergetics, we performed a Seahorse assay at 24 and 72 hours post treatment (**figure 7**). These results showed that pQ-EV-treated MCF7 spheroids have decreased oxygen consumption rate (OCR) levels 24 hours post treatment, indicating that pQ-EVs generate low OXPHOS levels (**figure 7a**). By contrast, pA-EV-treated spheroids have higher OCR and extracellular acidification rate (ECAR) levels relative to pQ-EV-treated spheroids, indicating that pA-EV treatment results in both high OXPHOS and high levels of glycolysis, likely driving the increased proliferation activity previously observed. UT spheroids produced OCR and ECAR levels between those of the two MSC-EV treatments. We next plotted the data of each section of the respiration profile, normalised to UT cultures (**figure 7b-e**). This analysis showed that basal respiration rates were not significantly affected by MSC-EV treatment (**figure 7b**). pA-EV treated spheroids experience higher levels of ATP-linked respiration, indicative of these cells’ increased energy demand in response to this treatment (**figure 7c**). Maximal respiration was also slightly higher for pA-EV-relative to pQ-EV-treated spheroids, indicating that pA-EV-treated spheroids have increased substrate availability and electron transport chain integrity (**figure 7d**). Finally, 24 hour pQ-EV treatments caused a significant decrease in spheroid spare capacity, compared to pA-EV and UT spheroids (**figure 7e**). Together, these results and the proliferation data indicate a more quiescent/glycolytic BCC phenotype with pQ-EV treatment.

At 72 hours, the energy map shows that the quiescent phenotype produced by pQ-EV treatment is less pronounced but indicative of a glycolytic metabolic bias (**figure 7f**). By contrast, at 72 hours, pA-EV-treated spheroids still exhibit a more energetic, aerobic phenotype compared to UT spheroids (**figure 7f-j**). The basal (**figure 7g**), ATP linked respiration (**figure 7h**) and spare capacity (**figure 7j**) of pQ-EV treated spheroids was still the lowest of all treatments, with pA-EV treated spheroids yielding comparatively higher levels of OXPHOS.

As stated in the introduction, our main interest concerns the metabolite cargo of the pQ-EVs produced from intact MSC monolayers and the pA-EVs produced from scratched MSC monolayers and to investigate whether these different cargos can elicit different dormancy and activation phenotypes in 3D cultured BCCs. Our results indicate that the two culture conditions do indeed produce different EV cargos and that these different cargos have different effects on the growth and respiration of MCF7 spheroid cultures. However, these cargoes are complex, as we illustrate above in our metabolomic and proteomic analysis. As such, we do not know which cargo components help to promote BCC dormancy or activation.

### pQ-EV cargo metabolites can influence MCF7 spheroid activity in isolation

Metabolites can both act as bioactive compounds and fuel biological processes and, as such, are a growing focus of research (23,24,73–75). We therefore assessed whether the pQ-EV metabolite cargo could cause a metabolic shift in MCF7 spheroids towards dormancy. To do so, we selected the MSC-EV cargo metabolites that showed the greatest difference between two MSC populations and used these to treat MCF7 spheroids. The resulting metabolite cocktail, called Group1, consisted of 40mM lactate+ 50mM glucose+ 8mM pyruvate, with rotenone administered in isolation as a positive control (blocking mitochondrial ATP production). Both Group 1 and rotenone were administered to MCF7 spheroids, which were then analysed for their cell proliferation and metabolism.

We performed an initial MTT assay to assess whether the chosen metabolite concentrations in Group 1 affected MCF7 viability for up to 72 hours of treatment. No significant effect on MCF7 viability was observed (**supplementary figure 8**). We then performed a colony formation assay, which showed that both Group1 and rotenone treatment caused significantly less colony formation relative to UT MCF7s (**figure 8a,b**), as was observed with the pQ-EV treatment. Both treatments also reduced MCF7 spheroid proliferation, as shown by Ki67 levels in western blots (**figure 8c-e**), as previously observed with the pQ-EV treatment. Propidium iodide analysis revealed that both treatments increased the proportion of MCF7 cells in G1 and decreased the proportion in S phase, compared to UT cells after 24 hours, particularly for Group 1 (**figure 8f**), again recapitulating the pQ-EV treatment result.

**Figure 8.**
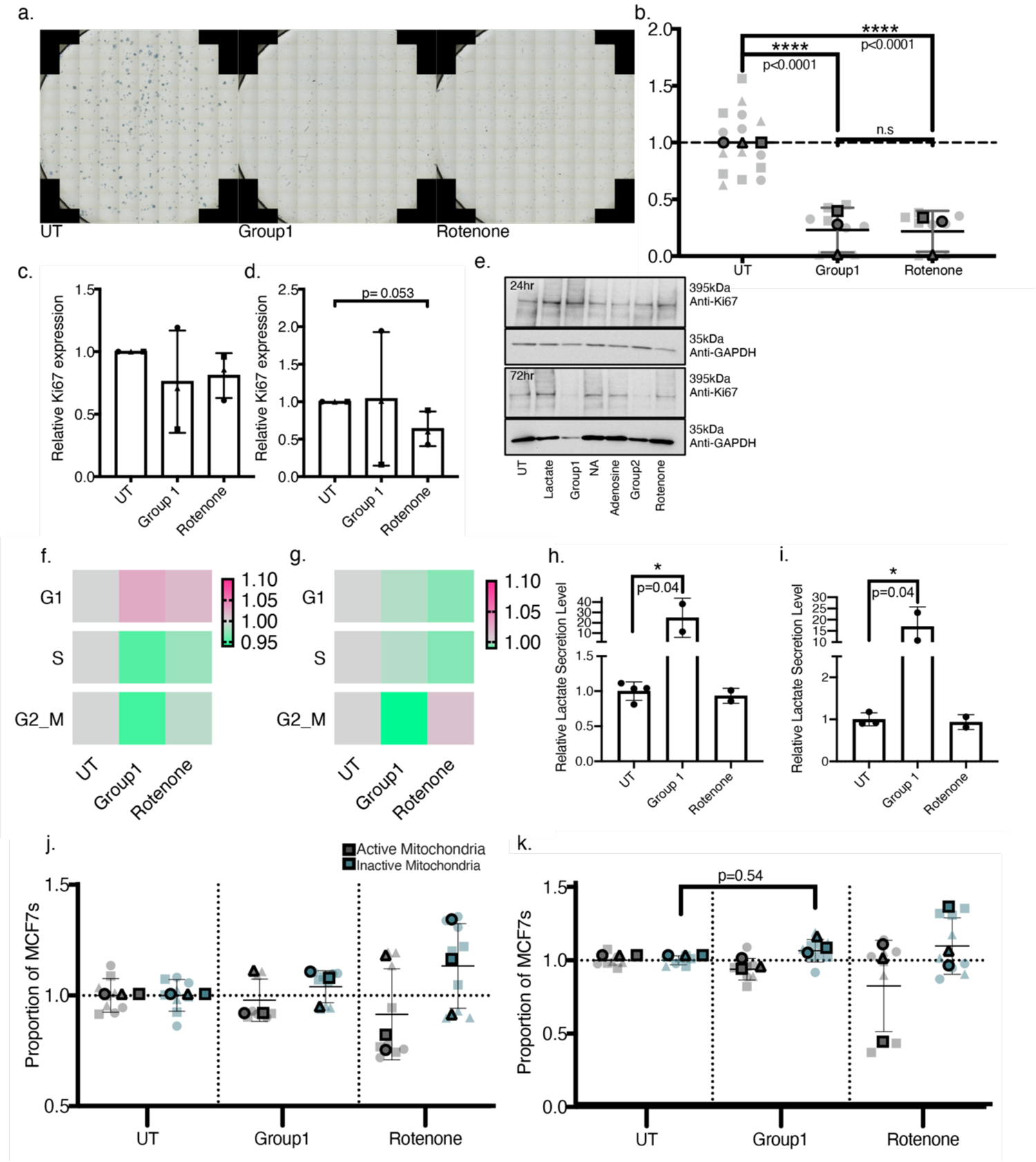
pQ-EV cargo metabolites can influence MCF7 spheroid activity in isolation. **a, b.**) Colony formation used assay to assess the effect of Group 1 metabolite mix on cultured MCF7 cells over 2 weeks. MCF7s were seeded at a low density in 2D monolayer. Cultures were either untreated (UT) or treated with DMEM-supplemented Group1 mix (40mM lactate+ 50mM glucose+ 8mM pyruvate) or 1µM Rotenone, every 3 days for 2 weeks, after which colonies were fixed, stained and counted. Treatments were normalised to the UT control. **a**.) Representative images**. b.)** Mean colonies per condition ± SD plotted, and an unpaired t-test performed, n=3. **c-e.)** 3D MCF7 cultures were UT or incubated for 24/ 72 hours with metabolite treatments before lysates were collected for Ki67 expression Western blot analysis. Ki67 band density was normalised to a GAPDH loading control and then further normalised to UT samples to calculate a fold-change value. **c.)** Quantification of Ki67 expression after 24 hour treatment. **d.**) Quantification of Ki67 expression after 72 hour treatment **e.)** Representative blot shown from n=3 independent blots. **(f&g)** 3D MCF7 spheroids were UT or incubated for 24/72 hours with metabolite treatments before being collected, dissociated with collagenase D, washed, fixed and stained with FxCycle™ PI/RNase Staining Solution (Life technologies; F10797). The stained single cell suspensions were analysed by flow cytometry. Graphs represent geometric mean fluorescent intensity per cell cycle phase, normalised to UT cells ± SD, with a minimum of 2000 cells analysed per 3 technical repeats. A 2-way ANOVA with multiple comparisons test was done, n=3. **f.**) Relative cell proportion per cell cycle phase after 24 hour treatment. **g.**) Relative cell proportion per cell cycle phase after 72 hour treatment**. (h,i.)** 3D MCF7 lactate secretion levels were measured to assess cellular levels of glycolytic metabolism following metabolite treatments. Supernatants were collected at each timepoint, and lactate levels quantified using the Lactate Glo^TM^ Assay (Promega, J5021). Graph depicts mean ± SEM concentration, n=2, with a minimum or 2 technical repeats run. **h.**) 24 hour timepoint. **i**.) 72 hour timepoint. **(j, k.)** 3D MCF7 cultures were untreated (UT) or incubated for **j**.)24 hours or **k.**) 72 hours with respective metabolite treatments and changes in mitochondrial respiration were evaluated using the JC-1 dye. Graphs depict mean ± SEM proportion of active and inactive mitochondria for each condition, normalised to UT cells. 3 technical repeats with a minimum of 2000 cells were measured per sample and an unpaired t-test completed between each condition, n=3.

We also observed that MCF7 lactate secretion (reflecting glycolysis) was increased at 72 hours following pQ-EV treatment and was dramatically increased in Group 1-treated MCF7 spheroids (**figure 8g**). Finally, metabolic analysis via JC-1 staining showed that both Group 1 and rotenone treatment decreased the levels of active mitochondria and increased levels of inactive mitochondria (**figure 8h**), again mirroring the pQ-EV treatment. These findings demonstrate that treating MCF7 cells with an exogenous metabolite cocktail that partially mimics the cargo of pQ-EVs can decrease the cells’ OXPHOS levels.

## Discussion

To advance our understanding of the process of BC dormancy, we need to better understand how the BM niche environment can induce and maintain dormancy by promoting cell cycle arrest and how it subsequently facilitates BC recurrence. It has been hypothesised that the BM provides a pro-dormancy signal via, amongst other cues, paracrine events that are regulated by resident MSCs (76). Recent evidence has also supported the involvement of MSC-EVs in the onset of BCC dormancy (3,13– 19,77,78). Here, we test the hypothesis that a change in MSC phenotype, caused by aging, trauma or acute therapeutic treatment, might induce BC recurrence. In this study, we therefore aimed to investigate whether MSCs secrete different EV cargoes following a change of phenotype in a manner that influences BCCs.

Our results show that MSC-EV cargos undergo distinct metabolomic and proteomic modifications when MSCs are placed under regenerative demand. We generated and investigated two distinct EV cargo populations: pQ-EVs (proposed quiescent, generated from intact MSC monolayers) and pA-EVs (proposed activating, generated from scratched MSC monolayers). Our findings show that the cargos of pQ-EVs contain a higher level of metabolites that are associated with glycolysis relative to pA-EVs. We observe in our metabolomic and proteomic screens that significant hits were more frequently observed in pQ-EVs. These results imply that pQ-EVs cargos contain a surplus of metabolites and proteins, including respiration-linked factors, which are depleted in the pA-EVs produced by MSCs under the energetic demand of regeneration. Moreover, our metabolomic pathway bioinformatic analyses linked these depleted metabolites and proteins in pA-EVs to the activation of pathways that are involved in the upregulation of cell growth and proliferation. Our proteomic analysis also identified protein-depletion patterns in EVs that could be linked to the upregulation of pathways involved in classic wound healing pathways in parent cells. Therefore, we have confirmed that a change in MSC phenotype, through the introduction of a regenerative demand, leads to the generation of different MSC-EV cargos in pQ-EVs and pA-EVs that appear to be linked to the activation status of the parent cell.

We next investigated whether pQ-EVs and pA-EVs could influence the behaviour of breast cancer cells in a 3D MCF7 spheroid model. As predicted from our bioinformatic analysis, pA-EVs induced an active MCF7 cell phenotype, with increased proliferation and migration relative to untreated or pQ-EV treated spheroids. From our metabolomic analysis, we also propose a novel mechanism for an MSC-EV-based metabolomic shift, in which pQ-EVs induce a shift towards MCF7 spheroid glycolysis, with classic hallmarks of dormancy, including a decrease in proliferation and migration. Conversely, pA-EVs support an MSC regenerative profile, more akin to dormancy escape or recurrence.

As discussed, bioactive metabolites are emerging as powerful signalling molecules that can drive cell processes (23, 73), as previously demonstrated in the control of stem cell differentiation (24,74,75). However, their role in cancer dormancy is not understood. Here, we provide the first evidence that in MSCs with low respiration (energy) demand, more metabolites are packaged into EV cargos and their delivery appears to reduce cancer cell growth. When MSCs are activated, fewer metabolites are packaged into EV cargos, and the lack of respiration-based metabolites in these cargos appears to create a demand for increased respiration (and cell activity) in the cancer cells. This data implies that regenerative demand placed on the bone of BC remission patients with dormant BCCs in their marrow, for example in response to traumatic injury or to elective procedures such as hip or knee arthroplasty where wound healing is initiated, could be activating factors in the exit from BCC dormancy and in secondary cancer formation.

We note that a major limitation in studying cancer dormancy is the lack of clinically relevant *in vivo* models. Although several transgenic models of spontaneous metastasis to bone exist, where cancer cells possess the ability to remain dormant before reactivation, these models often take months for disseminated tumour cells (DTCs) to become detectable and their metastasis reliability is weak (35, 79). While these transgenic models can recapitulate prolonged latency, the majority of in vivo studies rely on the inoculation of tumour cells directly into the animal’s circulation (80). These studies mimic bone colonisation without the entire metastatic cascade and often use highly aggressive cell lines that don’t show periods of dormancy (80–82). These limitations will need to be overcome for future *in vivo* MSC-EV/ BCC studies.

EVs have emerged as biological agents with substantial therapeutic potential, comprising a unique set of features that could allow them to be effective acellular drug delivery systems. For instance, can cross various biological tissue barriers, plasma membranes and endosomal membranes (83, 84). Moreover, endogenous cellular machinery can be used to produce the desired cargo, negating the challenges of manufacture, storage and loading of biotherapeutics (85). However, critically, there is a need for isolation processes to be safely standardised and massively upscaled, one of the main hurdles that will need to be overcome for EVs to be clinically successful (85). As we exemplify here, this EV scale up limitation can be overcome by exogenously developing the cargo identified as being therapeutically active. Our data allows us to envision a therapeutic strategy in which dormant BCCs are treated with a glycolytic metabolite cocktail, to ensure that these cells do not reactivate. Alternatively, treating dormant BCCs with glycolysis inhibitors or with an OXPHOS metabolite cocktail could ‘awaken’ the dormant BCC population, rendering them susceptible to chemotherapeutic treatments already developed for the clinic.

## Acknowledgements

Margaret Mullin for EM work. Ivana Milic for proteomics help. Glasgow Polyomics for running the Metabolics and RNA-sequencing experiments. Lauren Hope for running MSC marker flow panel.

## Authors’ contributions

SB: Responsible for the acquisition, analysis and interpretation of the data. Substantial contribution to the concept and design of the article. Approved the version to be published. YX: Assisted in the RNA sequencing analysis. Approved the version to be published. ER: Assisted in the acquisition of the proteomic data. Approved the version to be published. MJD: Substantial contribution to the data interpretation. Substantial contribution to the design and drafting of the article. Approved the version to be published. CCB: Responsible for the study conception and design of the work. Substantial contribution to data interpretation and the concept, design and drafting of the article. Approved the version to be published.

## Ethics approval and consent to participate

Not applicable.

## Consent for publication

Not applicable.

## DOI of raw data from the library

### Competing interests

The authors declare that there are no competing interests.

## Funding information

The authors received funding from Medical, Veterinary and Life Sciences DTP at the University of Glasgow and the Glasgow Polyomics Facility (ISSF Feasibility Study project funding) and BBSRC funded project (BB/L008661/1).

## Materials and Methods

### Cell culture

#### MSC culture

MSCs (Promocell©) were cultured using 10% FBS DMEM solution. MSCs were maintained in T75 tissue culture treated flasks and passaged until passage 4, to reach final 95% confluency. During each passage cells were washed using X1 PBS, removed following treatment with trypsin/versine solution, centrifuged (445g for 4 minutes) and reseeded into new flasks. Media was exchanged every 3 days.

#### MSC-EV collection

EV-free FBS (ThermoFisher, A2720803) was used instead of normal FBS to create collection media (CM) for EV isolation. CM was incubated with MSCs for 48 hours before collection.

#### MSC scratch wounding assay optimisation

MSCs were seeded at passage 4 in 10cm^2^ tissue culture treated petri dishes (ThermoFisher, 150464) at 1x 10^5^ cells/cm^2^ and grown until ∼95% confluent. A P200 tip was thereafter used to scratch across the dish, along a stripette placed above the dish, to achieve a straight scratch. The scratch pattern followed for each plate is detailed in Figure S1. Note, scratch injury healing time was determined using a time-lapse experiment, with images acquired every 20 minutes for 48 hours via the EVOS™ M7000 Imaging system (Figure S2).

#### MSC scratch injury assay for EV collection

pA-MSCs were scratched, washed twice with X1 PBS and thereafter incubated with 10mL of CM per dish for 48 hours. A minimum of 20 plates were set up per EV collection from pA-MSCs.

#### MCF7 2D culture

MCF7 2D cultures were generated as for MSCs, however used at passage 9-30 and maintained in Modified DMEM (DMEM containing 10%FBS and 20% Media 199).

#### Ultra-low attachment (ULA) method for 3D MCF7 spheroid formation

MCF7s were seeded on CellCarrier© Spheroid ULA 96-well Microplates (PerkinElmer, 6055330) at a density of 1x 10^4^ cells per well in Modified DMEM. Spheroids were formed after 24 hours and media exchanged every 3 days.

#### MCF7 culture MSC-EV treatment

3D MCF7 spheorid cultures were washed with X1 PBS and treated with fresh Modified DMEM supplemented with approximately 2×10^7^ MSC-EVs/ mL for 24 / 72 hours (unless otherwise stated). An untreated (UT) control was always included, which was treated with Modified DMEM only.

#### MCF7 culture metabolite treatment

3D MCF7 spheroid cultures were washed with X1 PBS and treated with fresh Modified DMEM supplemented with 1µM Rotenone, 40mM lactate, Group1 (40mM lactate + 50mM glucose + 8mM pyruvate), 100µM Adenosine, 10µM isonicotinic acid (NA) or Group2 (100µM Adenosine +10µM NA). An untreated (UT) control was always included, which was treated with Modified DMEM only.

#### Differential Ultracentrifugation for MSC-EV isolation

A minimum of 200mL CM was collected after being incubated with MSCs for 48 hours, per respective MSC condition. CM was then centrifuged in 50mL Falcon tubes for 20 minutes at 2000g, followed by 30 minutes at 10,000g to remove cell debris and any large apoptotic bodies. Thereafter, the resulting supernatant was transferred to 25mL ultracentrifuge tubes (Beckman Coulter, 355654) and centrifuged at 100’000g for 70 minutes using a fixed angle rotor, **Error! Reference source not found.**. T he supernatant was then discarded and EV pellets were re-suspended in 200μL in filtered X1 PBS. The 200μL resuspension was divided into 25μL aliquots. Aliquots were stored at −80°C and used immediately after thawing (86).

#### MSC-EV characterisation

MSC-EVs were characterised according to the Minimal information for studies of extracellular vesicles 2018 (MISEV 2018) using the following materials and methods (87).

#### MSC-EV quantification

The tuneable resistive pulse sensing (TRPS) technology (qNano IZON system; Izon, Cambridge, USA) was used to quantify MSC-EV concentration and size-distribution. The system was calibrated for voltage, stretch, pressure, and baseline current using the 100 and 200nm standard beads (Izon, CPC100 and CPC200). Thereafter, a dilution of MSC-EV and PBS suspension was run on a NP150 nanopore (for 70-420nm size range) and data analysis was performed by the qNano IZON software.

#### Dynamic Light Scattering (DLS)

Particle size distribution was analysed using Dynamic Light Scattering (DLS; Malvern Instruments, UK) and analysed using the Zetasizer software.

#### MSC-EV Transmission Electron Microscopy (TEM)

TEM was employed to assess the size, morphology and integrity of MSC-EVs. EV suspensions mixed with 2% paraformaldehyde (PFA)/PBS were placed on formvar-carbon coated EM grids. Grids were washed (X5, 10 minutes) in PBS and thereafter incubated in 1% glutaraldehyde. After 10 minutes, the grids are washed (×10, 2 minutes) in distilled H_2_O. A drop of uranyl oxalate solution (pH 7) was then transferred onto the grids and incubated at room temperature for 5 minutes. The grids were finally transferred to methylcellulose for 10 minutes on ice, dried and thereafter imaged with a JEOL 1200 Transmission Electron Microscope, with beam voltage of 80kV and a magnification of X6000. Tiff images were captured using Olympus Scandium Software.

#### Micro-BCA

The Pierce™ micro bicinchoninic acid (micro-BCA) reagent kit (ThermoFisher, 23235) was employed to measure the total protein content of isolated MSC-EVs. This kit was used according to the manufacturer’s guidelines, however the diluent used was 0.2M NaCl. Specifically, BSA standards were diluted in 0.2M NaCl to reach the respective standard concentrations and MSC-EVs were diluted 1:20 in 0.2M NaCl to reach 100µL.

#### 3D MCF7 culture MSC-EV uptake assay

To assess MSC-EV uptake into MCF7 cultures, EVs were stained using the PKH67 Green Fluorescent Cell Linker Mini Kit (Merk, MINI67-1KT) according to manufacturer’s guidelines. Unbound dye was removed through a 100’000g ultracentrifugation step and the dyed EV pellet resuspended in PBS. MCF7 spheroids were incubated with stained EVs in the dark for 4 and 24 hours, after which spheroids were fixed in 10% PFA, washed in X1 PBS, incubated with 200µM Höchst solution (ThermoFisher, 62249), washed twice in X1 PBS, and transferred to poly-D-lysine (Sigma, P7280) coated 8 well chamber slides. Spheroid EV uptake was visualised on the Inverted Nikon A1-R Spinning Disk Confocal system.

#### Live/Dead cell viability assay

LIVE/ DEAD^TM^ Viability/ Cytotoxicity Kit for mammalian cells (Invitrogen, L3224) was used to assess the viability of cell cultures. This kit enables live and dead cell discrimination, by simultaneously staining cells with green fluorescent calcein AM, demonstrating the intracellular esterase activity of live cells, as well as red-fluorescent ethidium homodimer, which indicates the loss of plasma membrane integrity exhibited by dead cells. Modified DMEM (MCF7s) or 10% DMEM (MSCs) containing 1μL mL^-1^ mix of calcein AM/ ethidium homodimer stain was added to cell cultures. After 30 minutes, the cells were washed and medium was exchanged for fresh, warm medium. Cells were then immediately visualised via the EVOS™ M7000 imaging system (TRITC channel; dead and FITC; alive). Images were processed in ImageJ.

#### MTT cell viability assay

3-(4,5-dimethylthiazol-2-yl)-2,5-diphenyltetrazolium bromide (MTT) is a yellow tetrazolium dye that is reduced to the insoluble formazan (purple) by oxidoreductase enzymes involved in metabolic processes in cells. The MTT assay was used to evaluate the metabolic activity of cells and as a representation of viability for proliferation. After 24/ 72 hour MCF7 MSC-EV treatments, MTT dye solution (5mg mL-1 MTT in PBS, pH 7.4) was added to each well of a 96-well plate and incubated for 2 hours. After the incubation, formazan crystals were solubilised with 200μL dimethyl sulphoxide (DMSO). The absorbance of each well was then read at 550nm on a Dynatech MR7000 microplate reader. Note, a blank sample (culture medium without cells) was used to calibrate the spectrophotometer to zero absorbance.

#### RNA extraction and isolation

RNA extractions of cell pellets were performed using the QIAGEN RNeasy mini kit, according to the manufacturer’s protocol. All centrifuge runs were done at 9000g for 15 seconds unless otherwise stated. Pellets were used either immediately after cell harvest or after having been stored at −80°C. 350µL of buffer RLT was added to each pellet and samples were homogenized by vigorous pipetting. 350µL of 70% ethanol was added to each lysate sample and mixed through inversion. The samples were directly transferred to Rneasy MinElute spin columns in 2mL collection tubes and centrifuged. The flow through in the collection tubes was then discarded. Subsequently, 350µL of buffer RW1 was added to the spin columns and centrifuged. 80µL of DNase 1 in buffer RDD was added directly onto the spin columns and incubated for 15 minutes at room temperature. Another 350µL of buffer RW1 was added to each spin column, after which they were centrifuged. The collection tubes were discarded and replaced, 500µL of buffer RPE added and the columns centrifuged again. 500µL of 80% ethanol was added and the columns centrifuged for 2 minutes. Subsequently, the collection tubes were discarded, spin columns placed in new collection tubes and centrifuged for 5 minutes at 13000g to dry the column membranes. The flow-through and collection tube were discarded, spin columns were placed in fresh 1.5mL collection tubes and 14µL of RNase-free water was added to the centre of each column membrane. The RNA was eluted by a 1 minute, 13000g centrifugation. RNA was quantified using a NanoDrop 2000 spectrophotometer (ThermoFisher Scientific, Waltham, MA, USA). The samples were then stored at −80°C.

#### Reverse Transcription (RT)

Reverse transcription (RT) was performed using a QuantiTect® Reverse Transcription Kit according to the manufacturer’s protocol. Template RNA samples were thawed on ice. The kit reagents (gDNA Wipeout buffer, Quantiscript® Reverse Transcriptase, Quantiscript® RT Buffer, RT Primer Mix, and RNase-free water) were thawed at room temperature. The solutions were gently mixed and centrifuged to collect any residual reagent clinging to the sides of the tubes. All ensuing reactions were prepared on ice. RNA content was normalised. The genomic DNA elimination reaction was set up with 2µL of gDNA Wipeout Buffer and the appropriate quantity of template RNA, made up to 14µL with RNase-free water. This reaction was incubated at 42°C for 2 minutes and then immediately transferred to ice. The reverse transcription reaction was set up with 1μL Quantiscript® Reverse Transcriptase, 4μL Quantiscript® RT buffer, 1μL RT primer mix, and 14μL template RNA from the genomic DNA elimination reaction, resulting in a total reaction volume of 20μL. This reaction mix was then incubated at 42°C for 15 minutes, followed by a 3 minute 95°C incubation, causing inactivation of the Quantiscript® Reverse Transcriptase. The reverse transcription reactions were stored at −20°C.

#### RT-qPCR

RT was performed using a QuantiTect® Reverse Transcription Kit according to the manufacturer’s protocol. The RT reactions were stored at −20oC. Using the primers detailed in Table, real-time RT-qPCR was performed based on SYBR Green and a 7500 sequence detection system.

#### Western blotting

Cell extracts were prepared in RIPA buffer (Tris-HCL pH 7.5 20mM, NaCl 150mM, EDTA 1mM, EGTA 1mM, NP40 1%, NaDOC 1%) containing protease inhibitor cocktail (Sigma, P8340) and protein content determined via the Pierce™ BCA Protein Assay Kit (ThermoFisher, 23225). Note, MSC-EVs were not lysed in RIPA buffer and protein content was determined via the Micro BCA™ Protein Assay Kit (ThermoFIsher, 23235).

Equal amounts of protein were loaded into Bolt™ 4 to 12%, Bis-Tris, 1.0 mm gels (ThermoFisher, NW04120BOX) and electrophoresis run in MOPS buffer (Invitrogen, NP0001). Protein was wet transferred to polyvinylidene difluoride (PVDF) membranes in transfer buffer (Invitrogen, NP0006). Membranes were blocked in 5% milk/ X1 PBS/0.1% Tween for 1 hour, followed by a 4°C overnight incubation in primary antibody diluted in 5% milk/ X1 PBS/ 0.1% Tween. Membranes were washed in X1 PBS/ 0.1% Tween and incubated in secondary HRP antibody for 1 hour at room temperature in 5% milk/ X1 PBS/ 0.1% Tween. HRP was visualised with ECL Western Blotting Substrate (Pierce™, 32106) and developed on a myECL™ imager (ThermoFisher). Note, to probe for Ki67 protein levels, cell lysates were run on NuPAGE^TM^ 3-8% Tris Acetate gels (Invitrogen EA0378BOX). To probe for CD63, non-reducing conditions were used.

#### Protein Profiler Array

The Proteome Profiler^TM^ Human XL Cytokine Array Kit (R&D Systems Inc) was employed to determine the relative levels of cytokines in the supernatants of pQ-MSCs and pA-MSC cultures and their respective EV populations.

All reagents were brought to room temperature before use and prepared according to the kits’ instructions. 2mL of Array buffer 6, which acts as a blocking buffer, was added to each well of the 4 well multi-dish. Each membrane was added to a separate well and incubated for an hour on a rocking platform. Thereafter, Array buffer 6 was aspirated and 1.5mL of sample was added to a well and incubated overnight at 4°C on a rocking platform. The next day each membrane was washed (X3, 10 minutes) in X1 Wash buffer and subsequently incubated with 1.5mL Detection Antibody cocktail. After a 1 hour incubation, membranes were again washed (X3, 10 minutes) in X1 Wash buffer and then incubated with 2mL of 1:2000 IRDye® 800CW Streptavidin (Li-Cor: 926-32230) in Array buffer 6 for 30 minutes. Thereafter, membranes underwent final washes (X3, 10 minutes) in X1 Wash buffer and were imaged on an Odyssey Sa infrared imaging system with Image Studio 4.0 software (Li-Cor).

To compare the different membranes the integrated density was measured for all cytokines, reference spots and negative control (background). The background was thereafter subtracted from each cytokine and the reference spots. All reference spots were averaged, and each cytokine was divided by the respective membrane reference spot average. Finally, the media only control was subtracted for each cytokine.

#### MSC supernatant preparation

MSCs were seeded at 1x 10^4^ cells/mL in 10% DMEM, and the proceeding day scratch wounding performed on half the samples. Supernatants were collected a 2, 24 and 48 hours post-scratch wounding timepoints. Supernatants were stored at −20°C until the assay was ready to be performed. A media only control sample was included here.

#### Colony Formation assay (CFA)

MCF7s were seeded at 150 cells/mL in a 12 well plate and treated with 2×10^7^ EVs/mL or respective metabolite treatments every 3 days for 2 weeks. MCF7s then washed in X1 PBS and stained/ fixed with 0.5% methylene blue/ 50% ethanol for 20 minutes at room temperature. Wells rinsed with distilled H_2_0, air-dried and imaged via the EVOS™ M7000 Imaging System.

#### 2D MCF7 Migration: µ-Slide chemotaxis

The effect of MSC-EV treatments on 2D MCF& migration was analysed using the Ibidi µ-Slide Chemotaxis kit, measuring directional motility of cells to treatments (two biological repeats). Following the manufacturers guidelines MCF7s were seeded at 3×10^5^ per mL, incubated for 4 hours and 60µL of the respective MSC-EV treatments or controls filled into the reservoirs. 24 hours post reservoir treatment, the slides were placed in the onstage incubator of the EVOS™ M7000 Imaging System and imaged on a X4 objective every 20 minutes for 24 hours. Results were analysed using the ImageJ plugin ‘Manual Tracking’ and Ibidi’s ‘Chemotaxis and Migration’ plugin tool.

#### 3D MCF7 Migration: Spheroid outgrowth measurements

The effect of MSC-EV treatments on 3D MCF7 spheroid size was investigated, facilitating the analysis of spheroid migratory projections and the overall proliferative effect of treatments. Upon 1, 3, 5 and 7 days of culture, spheroids were transferred to the EVOS™ M7000 Imaging system and images acquired. Respective treatments were repeated after every 72 hours. A minimum of 22 spheroids were imaged per treatment and the diameter of each spheroid was measured using ImageJ.

#### Flow cytometry

Flow cytometry buffer was used for all experiments, made up with X1 PBS supplemented with 0.5% BSA (SigmaAldrich, A9418) and 0.5mM EDTA (ThermoFisher, 15575020). All flow cytometry data analysis was carried out using the FlowJo software (BD).

#### 3D MCF7 spheroid dissociation for flow cytometry analysis

MCF7 spheroids were generated and treated with the respective treatment. 24/ 72 hours after treatments, spheroids were washed with X1 PBS and 10 spheroids pooled per sample, with three repeats per sample. To dissociate the spheroids, 2.5mg/mL Collagenase D (Merk, 11088858001) /PBS was filter sterilized and added to spheroids for 30 minutes at 37°C. The spheroids were dissociated by carefully pipetting up and down after 15 minutes and again after the entire 30 minute incubation.

#### MSC stemness/ differentiation analysis

The analysis of MSC stemness was carried out using a flow cytometry antibody panel (Table 2). MSCs were seeded at 1x 10^4^ cells/mL in 10% DMEM, and the proceeding day the scratch assay performed on half the samples. 2, 48 and 120 hours after treatment, cells were washed with X1 PBS, harvested with TrypLE (Gibco^TM^, 12604013), washed twice in X1 PBS and incubated with the respective antibody cocktail for 1 hour at 4°C with occasional agitation. Unstained, full stain and fluorescence minus one (FMO) controls were included, using unscratched MSCs. Following incubation, cells were washed in flow cytometry buffer and resuspended in 200µL before running on the cytometer (samples were prepared by me, however due to COVID-19, the samples were kindly run on the flow cytometer by Lauren Hope).

**Table 2.** Primary Western Blot Antibodies used.

| Primary Ab | Dilution | Cat No |
| --- | --- | --- |
| GAPDH | 1: 1000 | Abcam, ab8245 |
| CD63 | 1: 2000 | Abcam, ab59479 |
| Annexin V | 1:2000 | Abcam, ab14196 |
| Calnexin | 1: 4000 | Abcam, ab10286 |
| Ki67 | 1: 4000 | Abcam, ab92742 |

**Table 3.**
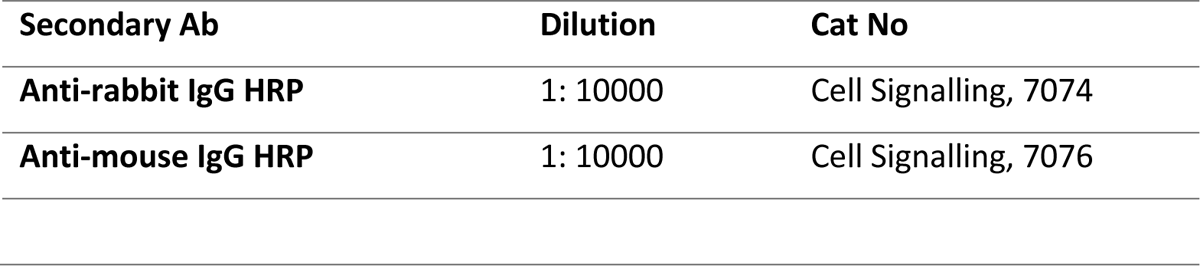
Secondary Western Blot Antibodies used.

**Table 0.** MSC stemness/ differentiation antibody panel

| Antibody | Conjugate | Volume per condition (µL) |
| --- | --- | --- |
| Lin 1 | FITC | 3 |
| CD34 | APC | 1.5 |
| CD73 | PE | 3 |
| CD45 | Pacific blue | 1.5 |
| CD90 | PE-Cy7 | 3 |
| CD105 | PerCP-Cy5.5 | 3 |

#### Propidium Iodide (PI) cell cycle analysis

For analysis of cellular proliferation by PI incorporation, the FxCycle™ PI/RNase Staining Solution (Life technologies; F10797) was used. MCF7s were seeded at 1×10^4^ cells/mL and treated with both pQ-EV and PA-EV at 2×10^7^ MSC-EVs/mL the next day. 24/ 72 hours after EV treatments, cells were washed with X1 PBS and harvested with TrypLE (Gibco^TM^, 12604013). Thereafter, cells were dissociated, washed twice with X1 PBS and fixed by adding 70% ethanol to the cell pellet, dropwise whilst vortexing, and stored at 4°C for 30 minutes. Once fixed, the ethanol-cell suspension was spun down at 1500g for 5 minutes and the resulting cell pellet washed in X1 PBS. The washed cell pellets were resuspended in 500µL FxCycle™ PI/RNase Staining Solution (Life technologies; F10797) and analysed via Attune NXT flow cytometer (ThermoFisher).

#### JC-1 mitochondrial activity assay

Mitochondrial function was quantified via the membrane potential dye JC-1 (ThermoFisher, M34152). MCF7s and MSCs were cultured in basal media containing 2µM JC-1 for 30 minutes at 37°C. 2D cultures were thereafter detached with TrypLE (Gibco^TM^, 12604013), with 3D cultures being dissociated as previously described and analysed by flow cytometry. As a positive control for mitochondrial depolarisation, cells were treated with JC-1 as well as 50µM carbonyl cyanide 3-chlorophenylhydrazone (CCCP; SigmaAldrich, C2759).

#### Cellular Glucose Uptake

Cellular glucose uptake was measured by culturing cells in glucose free DMEM for 2 hours before adding 100µM of the fluorescent glucose analogue 2-(N-(7-Nitrobenz-2-oxa-1,3-diazol-4-yl) Amino)-2-Deoxyglucose (2-NBDG; ThermoFisher, N13195). The analogue was incubated with MCF7s for 30 minutes and MSCs for 60 minutes at 37°C, upon which cells were washed in X1 PBS. 2D cultures were thereafter detached with TrypLE (Gibco^TM^, 12604013), with 3D cultures being dissociated as previously described and fluorescence incorporation measured analysed by flow cytometry.

#### Lactate secretion measurement

To measure the level of L-Lactate secretion by MSCs/ MCF7s the Lactate-Glo^TM^ Assay (Promega, J5021) was employed. This bioluminescent assay couples lactate oxidation and NADH production with a bioluminescent NADH detection system. The luminescent signal produced is proportional to the amount of lactate in the sample and increases until all the lactate is consumed.

The lactate detection reagent was prepared according to the kit’s instructions. Cells were incubated with their respective treatments, made up using 10% dialysed FBS (A3382001, ThermoFisher) in DMEM. Prior to running the full assay, a dilution factor had to be determined to ensure that the levels of lactate were in the detectable range of the kit. All samples were diluted ×10’000 in X1 PBS, except 3D MCF7 supernatants which were subject to a ×1000 dilution in X1 PBS. 50µL of sample was added to a white bottom 96-well assay plate and subsequently 50µL of freshly made lactate detection reagent added. The plate was shaken for 30 seconds and incubated for 60 minutes at room temperature. Luminescence was recorded using a Pherastar FS plate reader (BMG Labtech). A standard curve allowed for the concentration of L-Lactate in the samples to be determined.

#### Seahorse mitochondrial bioenergetics assay

Oxygen consumption rate (OCR) and extracellular acidification rate (ECAR) were measured using the Agilent Seahorse XF24 Analyser (2D MCF7 cultures) and Agilent Seahorse XFe96 Analyser (3D MCF7 cultures). Mitochondrial respiration was determined by exposing MCF7s treated with MSC-EVs to mitochondrial perturbing agents Oligomycin (10μM, Sigma), FCCP (6μM, Sigma) and rotenone plus antimycin A (1μM, Sigma).

**Table 3.** Seahorse media recipe

| Reagent | Component | Cat No | Final Concentration |
| --- | --- | --- | --- |
| **XF Seahorse media** | XF base medium | Agilent, 103334-100 |  |
|  | Pyruvate | Agilent, 103578-100 | 2mM |
|  | Glucose | Aglient, 103577-100 | 10mM |
|  | L-Glutamine | Aglient, 103579-100 | 2mM |

MSC-EV treatment basal respiration relative to UT was calculated as the average of the last measurement rate before oligomycin injection minus average of non-mitochondrial respiration rate. MSC-EV treatment maximal respiration rate relative to UT was calculated as the maximum rate measurement average after FCCP injection, minus the average of non-mitochondrial respiration. ATP-linked relative to UT was calculated as average of maximum rate measurements before oligomycin injection minus the average of minimum rate measurements after oligomycin injection. Spare capacity indicates the difference between the average maximal respiration rate ATP-linked respiration, proton leak levels and non-mitochondrial respiration.

#### 3D Seahorse mitochondrial bioenergetics assay

For spheroid culture mitochondrial bioenergetics analysis, MCF7 spheroids were generated and treated respectively. 100μg/mL Poly-D Lysine (Sigma, P7280) was used to coat Agilent Seahorse XFe96 Spheroid Microplates (Agilent, 102905-100), which were incubated with XF media (Table) in a non-CO_2_ 37°C incubator 1 hour prior to use.

Once MSC-EV incubation timepoints were reached, spheroids were transferred to the prepared Agilent Seahorse XFe96 Spheroid Microplates and these then placed in a non-CO_2_ incubator for 1 hour prior to running the experiment on the XFe96 analyser. Data analysis was performed via the Agilent’s Seahorse Analytics software.

#### MSC-EV Metabolomics screen

Metabolomic analysis was performed on whole isolated MSC-EVs. Three separate repeats of pQ-EV and pA-EV isolations were pooled upon normalization for particles/mL for ever sample and used for hydrophilic interaction liquid chromatography-mass spectrometry analysis (UltiMate 3000 RSLC, ThermoFisher, with a 150 x 4.6mm ZIC-pHILIC column running at 300µL/min^-1^ and Orbitrap Exactive). This was provided as a service by Glasgow Polyomics Facility. A standard pipeline, consisting of XCMS (peak picking), MzMatch (filtering and grouping) and the Glasgow Polyomics PiMP software (additional filtering, post-processing and identification) was used to process the raw mass spectrometry data. Identified metabolites were validated against a panel of unambiguous standards by mass and predicted retention times. Heatmaps of metabolites of interest were generated using the MetaboAnalyst software (version 5.0). Pathway analysis was performed using Qiagen’s Ingenuity Pathway Analysis software.

#### MSC-EV Proteomics screen

Mass spectrometry bottom-up and label free proteomic analyses was performed on MSC-EVs to uncover their proteomic cargo.

For this protocol, the EVs were isolated using size exclusion chromatography (Izon qEVoriginal). The isolated EVs were then further concentrated using 0.5mL 3kDa Amicon spin columns. EV protein concentration was quantified using a micro-BCA assay (0). Samples were thereafter sent to Dr Ewan Ross at Aston University, Birmingham, where with the help of his colleague Dr Ivana Milic, the remainder of this protocol was carried out.

Firstly, the samples were reduced in Laemmli buffer (for 15 minutes at 65°C) and then loaded and separated on 10% SDS-PAGE gel. The resolved proteins were stained with Coomassie G250 blue overnight at 4°C. After destaining the gel, all samples were divided into 5 bands across the whole gel and separately excised and further destained in 50% acetonitrile in 50mM ammonium bicarbonate. Destained gel sections were then dehydrated and vacuum dried after which they were rehydrated for proteolytic in-gel digestion. The following morning, extraction of peptides from gel sections was done sequentially in 30% and 50% of acetonitrile in 50mM ammonium bicarbonate (15 minutes in an ultrasonic bath), and subsequently samples were fully dehydrated in pure acetonitrile. With each dehydration step, peptide extracts from a single sample section were combined into one polypropylene tube, after which they were vacuum dried and stored at −20°C prior to mass spectrometry analysis.

These samples then underwent liquid chromatography-coupled tandem mass spectrometry (LC-MS/MS) analysis (nanoEase M/Z Symmetry C18 Trap Column, 100Å, 5 µm, 180 µm x 20mm, using a nUPLC system; flow rate of 5µL/min of eluent B (eluent B: acetonitrile in aqueous 0.1% formic acid)). Stable peptide electrospray was formed at 2500V using a PicoTipTM emitter, which allowed for the charged peptides to be infused into 5600 TripleTof mass spectrometer (AB Sciex). Relative protein quantification was performed using Progenesis QI for proteomics software (version 4, Nonlinear Dynamics, UK). Data was further in-silico normalised (based on the peptide distribution) to allow for high-accuracy relative quantification.

This data was then analysed by me using MetaboAnalyst software (version 5.0) and Qiagen’s Ingenuity Pathway Analysis software.

#### RNA-seq analysis of MSC-EV treated spheroids

MCF7 spheroids were generated and treated for 24/72 hours before 20 spheroids were pooled per sample, washed twice in X1 PBS and spheroid pellets lysed using Qiagen RLT lysis buffer. RNA isolation was performed as previously described.

Sequencing libraries were then prepared from total RNA using the Illumina TruSeq Stranded mRNA Sample Preparation Kit. Libraries were sequenced in 75 base, paired end mode on the Illumina NextSeq 500 platform. Raw sequence reads were trimmed for contaminating sequence adapters and poor-quality bases using the program Cutadapt (88). Bases with an average Phred score lower than 15 were trimmed. Reads that were trimmed to less than 54 bases were discarded. The quality of the reads was checked using the Fastqc program (http://www.bioinformatics.babraham.ac.uk/projects/fastqc/) before and after trimming. The reads were “pseudo aligned” to the transcriptome using the program Kallisto (89). The differential expression for the analysis groups were assessed using the Bioconductor package DESeq2 (90). This was provided as a service by Glasgow Polyomics Facility. Further analysis and figure generation was performed via Cluster and Metascape resources.

## Supplementary Information

**SFigure 1.**
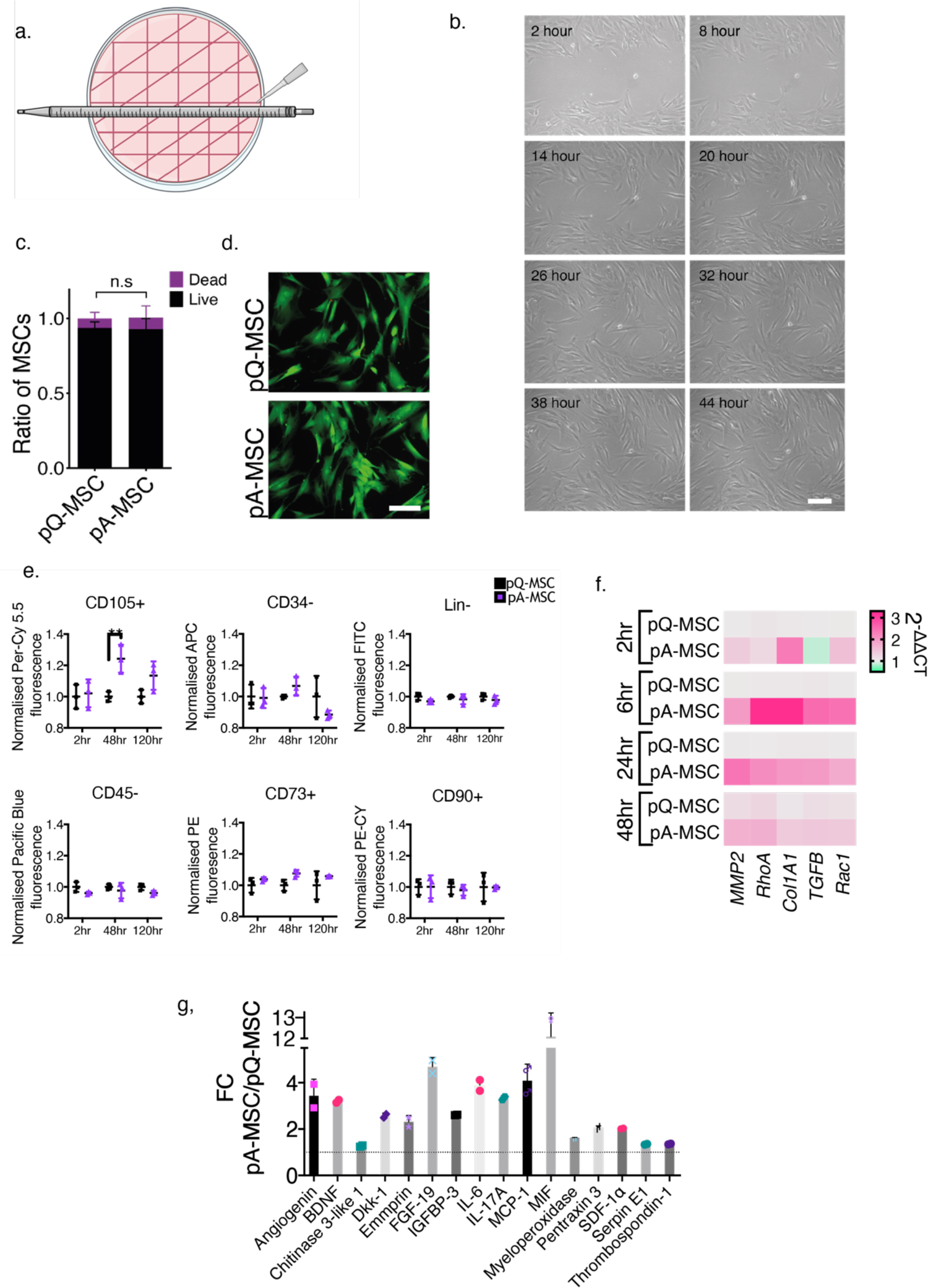
Development of a regenerative MSC model. **a.**) The MSC scratch assay was performed using a P200 tip along a stripette as illustrated, to maintain consistency across experiments. **b.**) A time-lapse imaging experiment was performed to assess the time period required for an entire scratch wound to heal. Scale bar represents 100 µm, n=1. **c, d.**) MSC cultures were untreated (pQ-MSC) or scratch-injured (pA-MSC) and incubated with EV collection media (CM). The LIVE/ DEAD^TM^ Viability/ Cytotoxicity Kit for mammalian cells (Invitrogen, L3224) was used to investigate the viability of these MSCs after a 48 hour CM incubation. An unpaired t-test was performed between samples, n=3. **d**.) Representative images. Scale bar represents 100µm. **e.**) MSC cultures were untreated (pQ-MSC) or scratch-injured (pA-MSC) and cultured for 2, 48 or 120 hours before collection for flow cytometry, being run against a panel of markers associated with MSC stemness (see Table). Here the marker-conjugate fluorescent intensity was assessed to identify the state of pQ-MSCs and pA-MSCs stemness over time. Graphs show the mean fluorescent intensity ± SD of 3 technical replicates for each MSC population, normalised to mean pQ-MSC intensity per timepoint. A 2-way ANOVA test with multiple comparisons performed, **= p< 0.01 (p=0.0051), n=1. **f.**) MSC cultures were untreated (pQ-MSC) or scratch-injured (pA-MSC) and cultured for 2, 48 or 120 hours before collection for flow cytometry, being run against a panel of markers associated with MSC stemness (see Table). Here the marker-conjugate fluorescent intensity was assessed to identify the state of pQ-MSCs and pA-MSCs stemness over time. Graphs show the mean fluorescent intensity ± SD of 3 technical replicates for each MSC population, normalised to mean pQ-MSC intensity per timepoint. A 2-way ANOVA test1 with multiple comparisons performed, **= p< 0.01 (p=0.0051), n=1. **g.**) MSC cultures were untreated (pQ-MSC) or scratch-injured (pA-MSC) and cultured for 48 hours before collection of RNA for RT-qPCR analysis. The migration and cellular activity markers assessed demonstrated a clear trend, with pA-MSC gene expression upregulated for most genes and timepoints, compared to pQ-MSCs. Expression data for each gene was normalised to a *GAPDH* housekeeping control and then normalised to pQ-MSCs to generate fold-change values. Heatmap depicts mean values from n=3 experiments. Upregulated genes are in pink and downregulated genes are in green. One-way ANOVA performed followed by Kruskal-Wallis test with multiple comparisons, no significant differences detected. **h.**) MSC cultures were untreated (pQ-MSC) or scratch-injured (pA-MSC) and cultured for 48 hours before collection of MSC supernatant for analysis. The Proteome Profiler^TM^ array (R&D Systems) was employed for parallel determination of relative levels of secreted cytokines from pQ-MSC and pA-MSC cultures. Graph depicts mean of 2 technical repeats ± SD and cytokine abundance normalised to pQ-MSC levels. Hits presented demonstrated a minimum significant fold-change (FC) of **= p<0.01 between the two samples, calculated via a 2-way ANOVA with multiple comparisons test, n=1.

**Table S1.** MSC Stemness Markers, Antibodies and Conjugates

| Positive markers |  | Negative markers |  |
| --- | --- | --- | --- |
| Antibody | Conjugate | Antibody | Conjugate |
| CD73+ | PE | Lin- | FITC |
| CD90+ | PE-CY7 | CD34- | APC |
| CD105+ | PerCP-Cy 5.5 | CD45- | Pacific Blue |

**SFigure 2.**
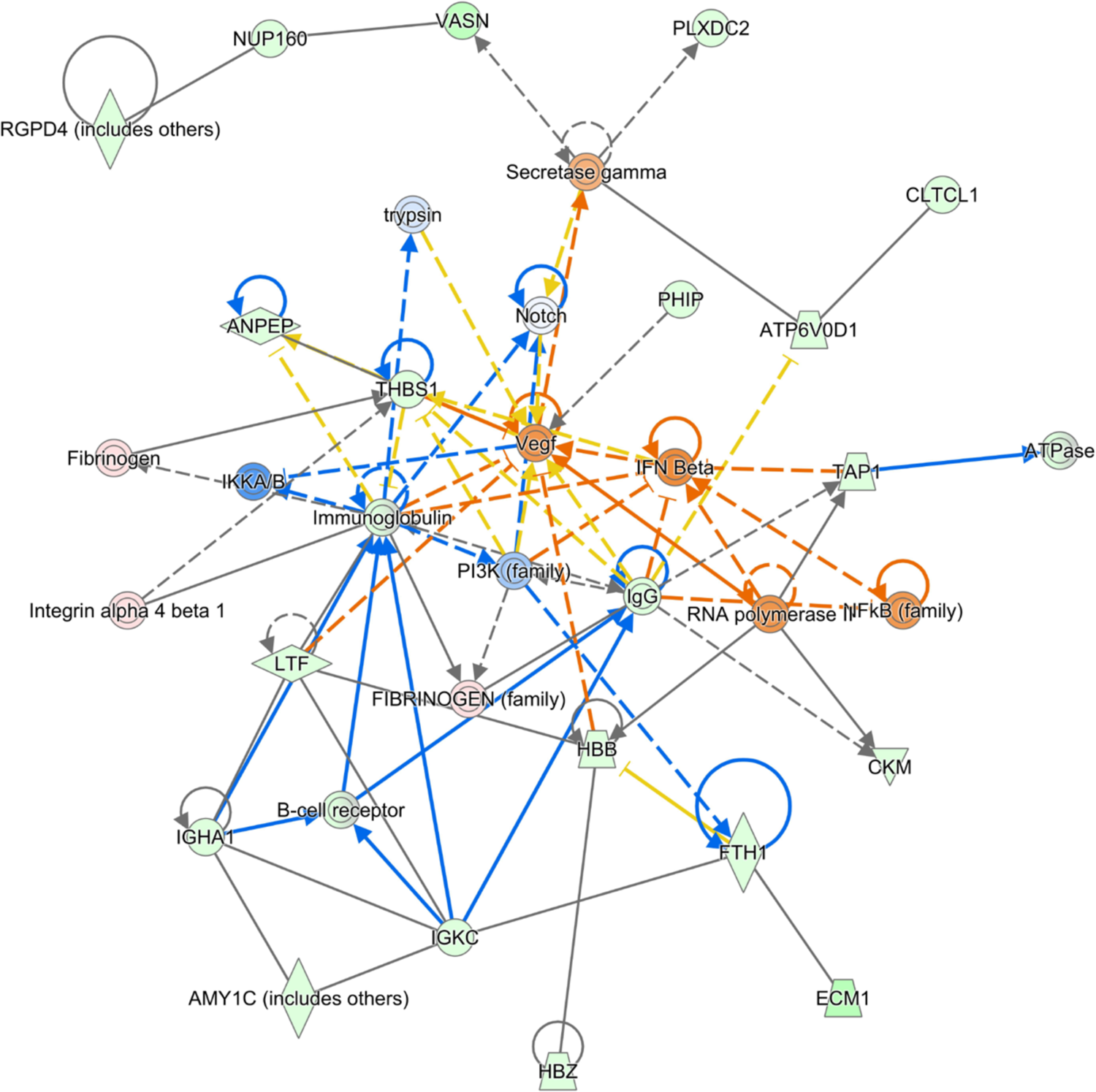
Proteomics screen reveals differences in MSC-EV cargo, with pA-EV cargo linked to pathological pathways. The proteomic cargo of pQ-EV and pA-EV samples was uncovered through liquid chromatography-coupled tandem mass spectrometry (LC-MS/MS) analysis, n=3. Normalised peak intensities fold-change values of pA-EV vs pQ-EV were subject to a Comparison Analysis through Ingenuity pathway analysis (IPA, Qiagen) and the top 2 network pathways mapped, n=3. Graph presents second mapped pathway which depicts the predicted activation of growth and inflammatory response pathways.

**SFigure 3.**
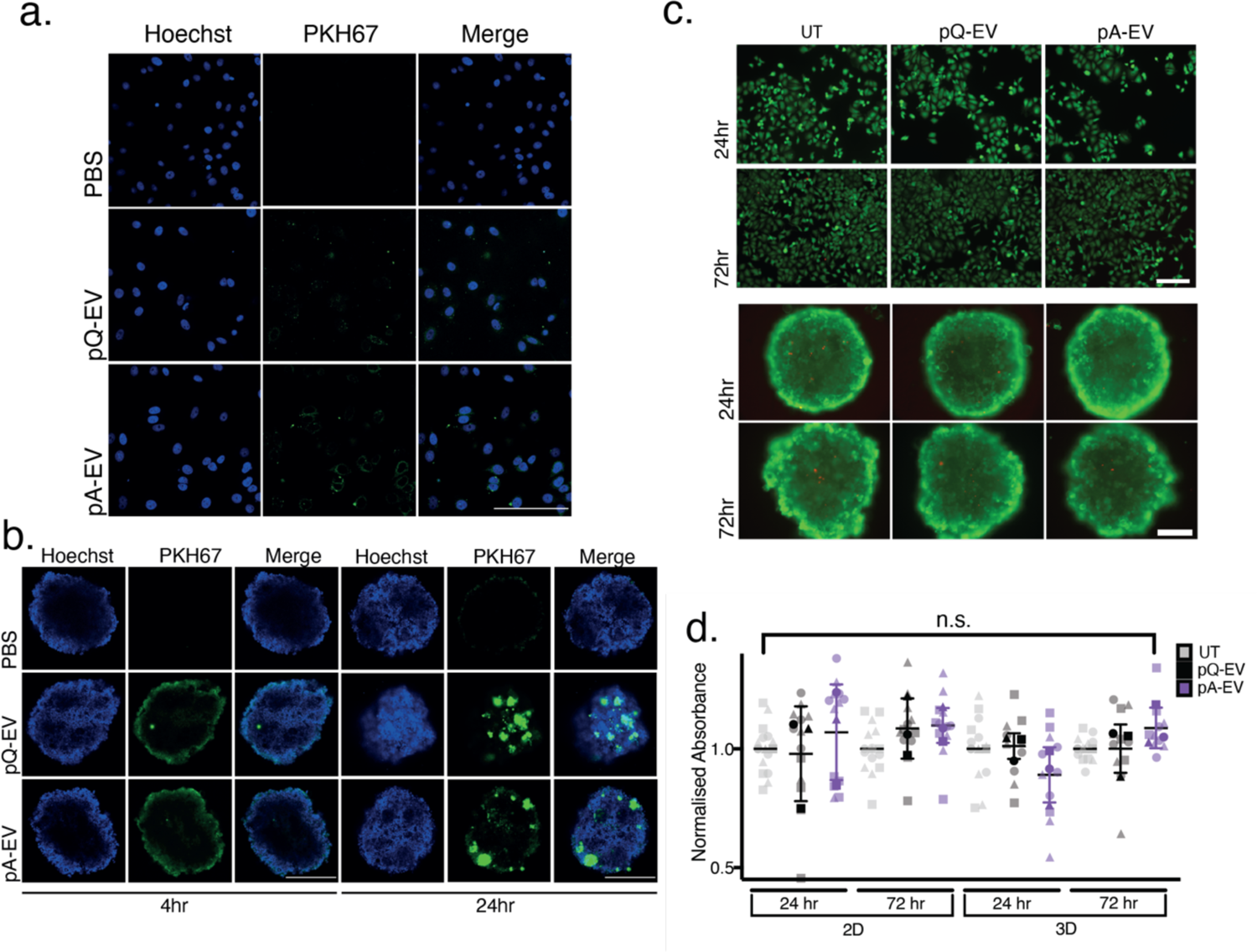
MCF7 cultures can successfully uptake MSC-EVs without influencing cell viability 2D and 3D MCF7 cultures were untreated (UT) or incubated with isolated and characterised pQ-EV and pA-EV treatments, assessing whether cultures could incorporate EVs without influencing cell viability. **a**.) Representative confocal microscopy analysis images of PKH67 stained MSC-EVs incubated with 2D MCF7s for 4 hours. A stained PBS control was used. Nuclei were stained with Hoechst and scale bare represents 500µm. **b**.) Representative confocal microscopy analysis images of PKH67 stained MSC-EVs incubated with 3D MCF7s for 4 and 24 hours. A stained PBS control was used. Nuclei were stained with Hoechst and scale bare represents 500µm. **c**.) Representative images from LIVE/ DEAD^TM^ Viability/ Cytotoxicity Kit for mammalian cells (Invitrogen, L3224), performed to further assess the viability of MCF7 cultures upon MSC-EV treatment. Here calcein AM staining reveals the presence of live cells (green), with ethidium homodimer staining dead cells (red). Scale bar represents 100µm. **d**.) Cell viability was observed after MTT treatment upon 24/ 72 hours of MCF7 spheroid MSC-EV treatments. Graph represents mean absorbance normalised to UT cells ± SEM. A 2-way ANOVA with multiple comparisons showed no significant difference between samples, n=3.

**SFigure 4.**
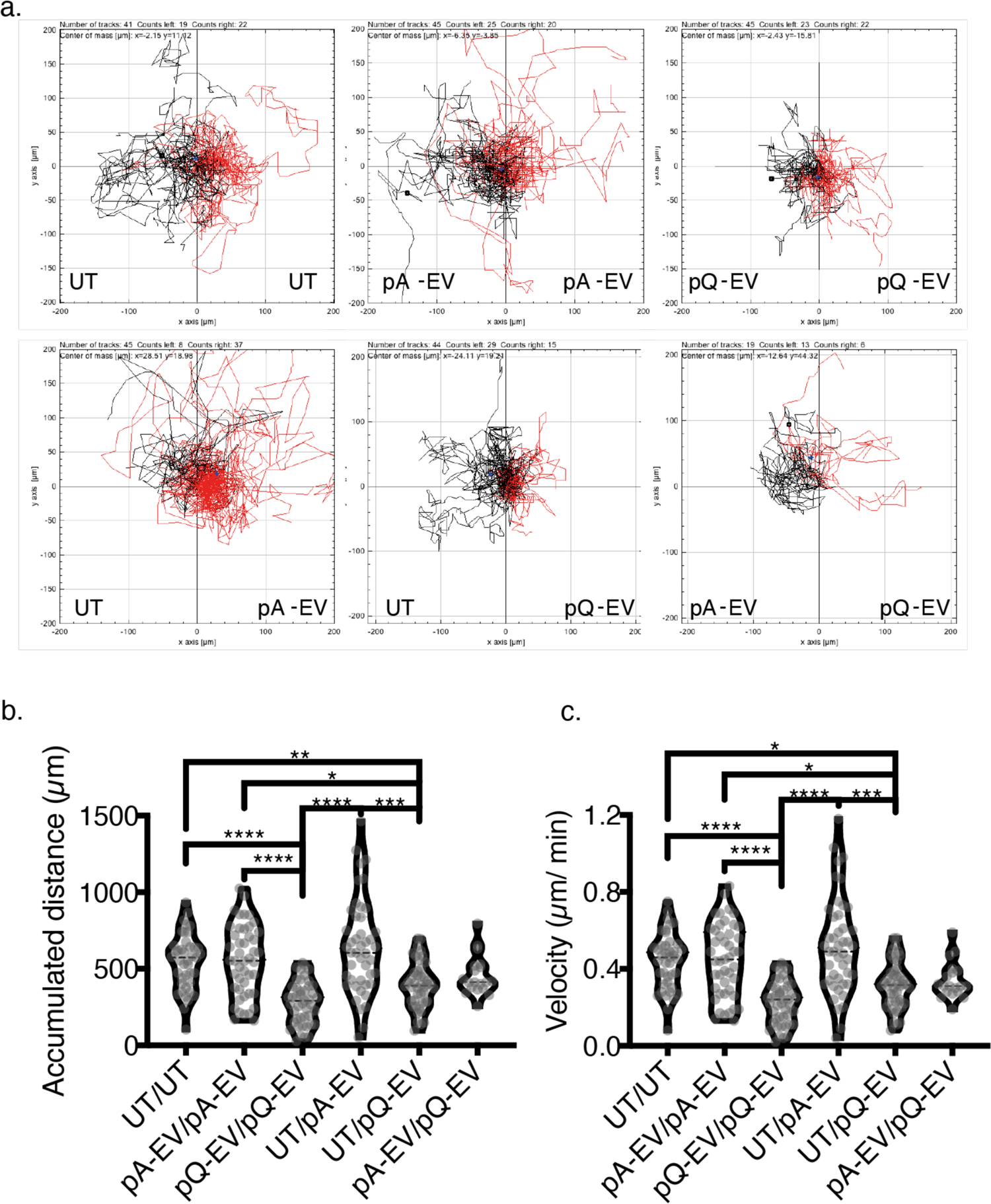
The effect of MSC-EV treatments on 2D MCF7 migration. The effect of MSC-EV treatments on 2D MCF7 migration was investigated via single cell tracking analysis. This was facilitated though Ibidi chemotaxis slides, demonstrating chemoattractant/ repellent effects of treatments. Upon 24 hours untreated (UT), CMSC-EV and SMSC-EV treated slides were transferred to the EVOS M7000 Imaging system and images acquired every 20 minutes to create an image sequence capturing the 24-48 hour treatment period. A minimum of 24 cells were tracked per treatment using the Manual Tracking ImageJ plugin and the migration of the cells was analysed via the Chemotaxis Tool ImageJ plugin. **A.**) Wind-rose plots show cell tracks for each condition, n=2 independent experiments combined. **B.**) Violin plot shows mean accumulated distance per tracked cell, n=2. **C.**) Violin plot shows mean velocity per cell tracked, n=2. B, C.) One-way ANOVA followed by a Kruskal Wallis multiple comparisons test was done for accumulated distance and velocity measurements, *= p< 0.05, **= p< 0.01, ****= p< 0.0001.

**SFigure 5.**
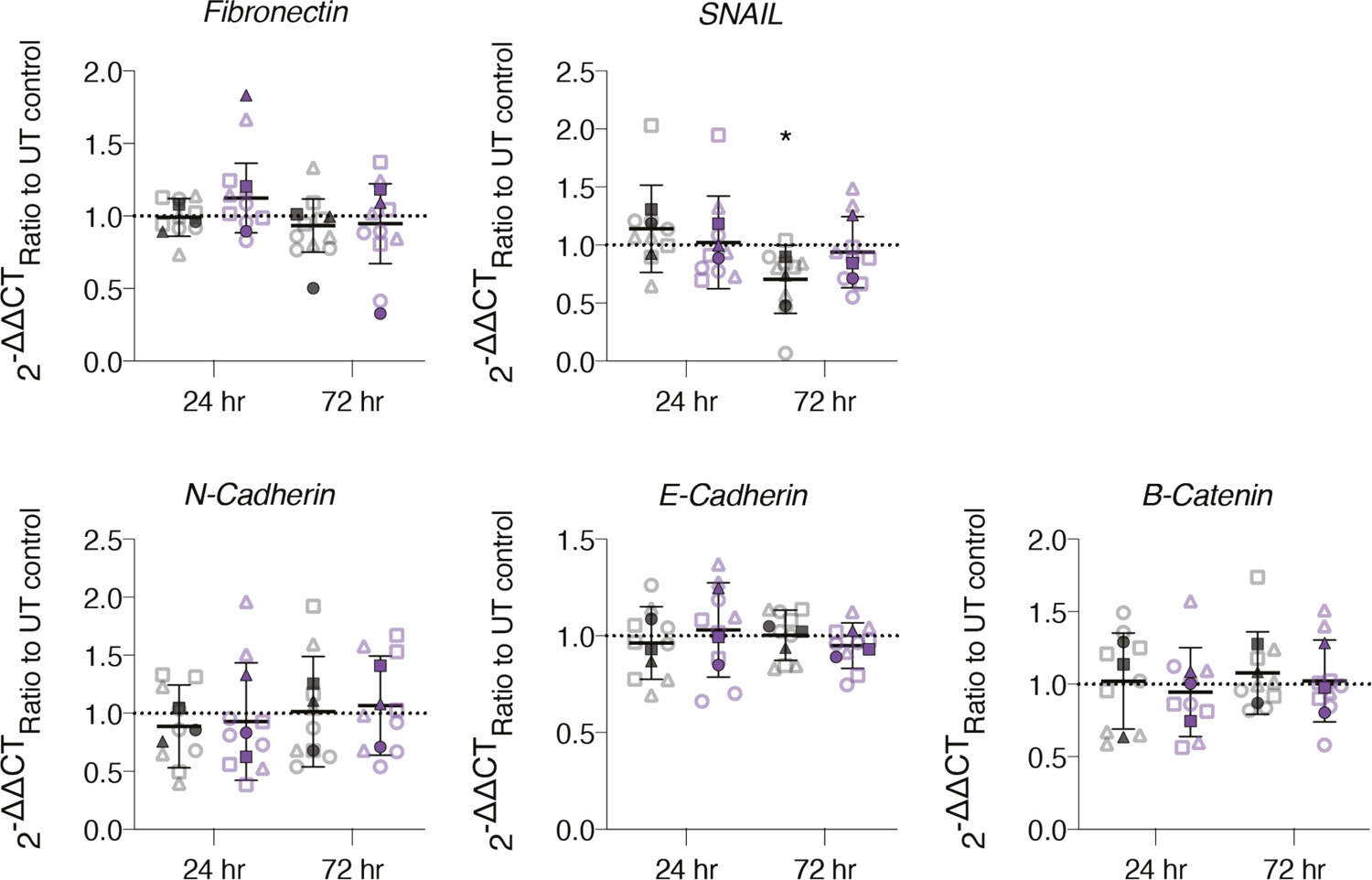
Effect of MSC-EV treatments on 3D MCF7 spheroid EMT marker expression 3D MCF7 cultures were UT or incubated for 24/ 72 hours with pQ-EV/ pA-EV treatments before RNA extraction for RT-qPCR analysis. Expression data for each gene was normalised to *GAPDH* housekeeping gene, then normalised to UT cells to generate 2^-ΔΔCT^ values. Mean ± SD plotted for each condition with 3 technical repeats done for each independent experiment. A one-way ANOVA followed by a Kruskal Wallis multiple comparisons test performed with no significance found, n=3.

**SFigure 6.**
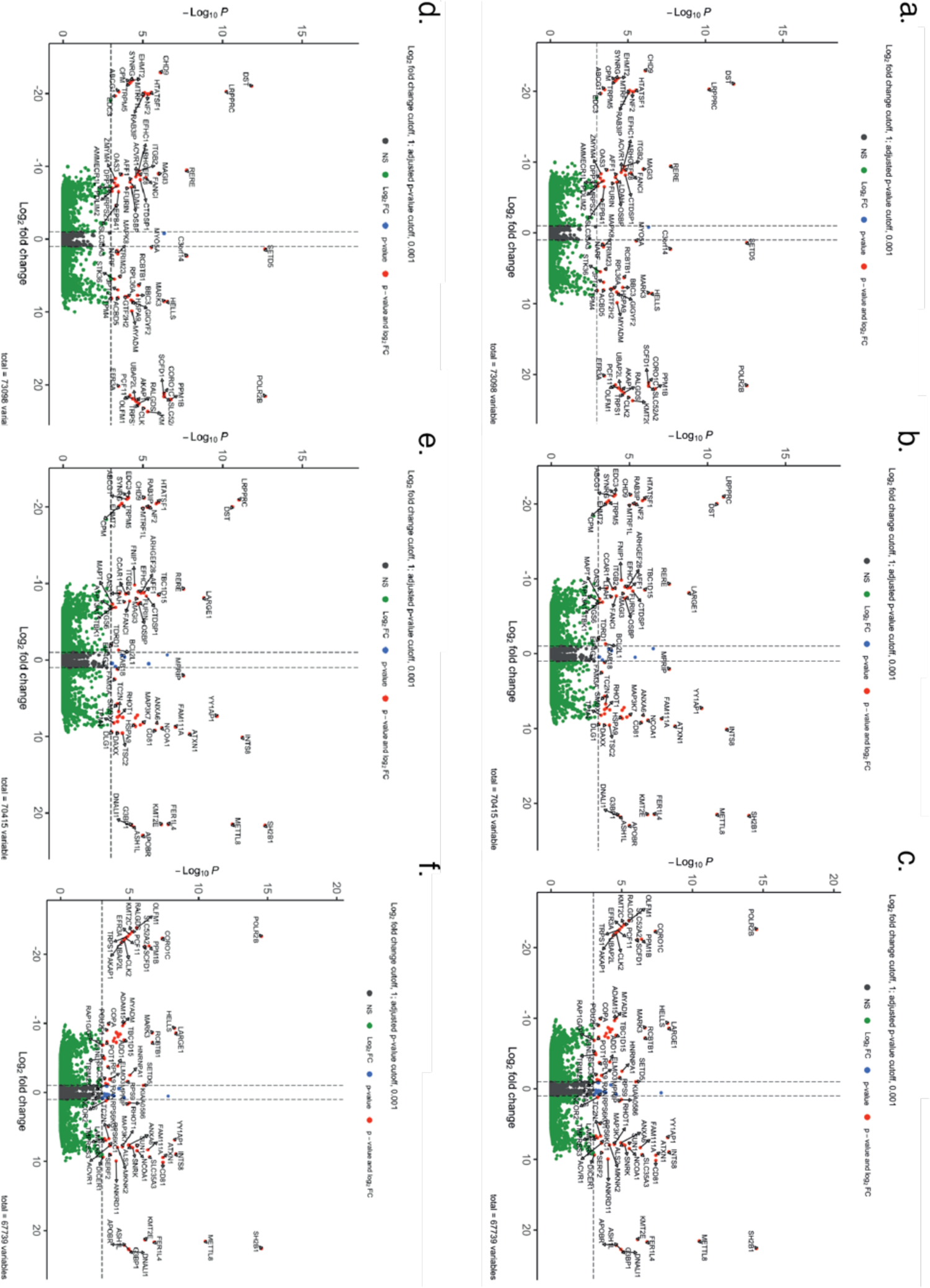
RNA-seq volcano plots. **a-f**.) Volcano plots show all hits per condition, with fold-change (FC) plotted against p-values. Grey coloured points represent non-significant (NS) hits, green points are Log2 fold-change (FC) hits and red points represent hits with Log2FC and where p<0.001. **a.**) 24 hour pQ-EV vs pA-EV, **b.**) 24 hour pQ-EV vs UT, **c.**) 24 hour pA-EV vs UT. **d-f**.) Represent the 72 hour treatment conditions.

**SFigure 7.**
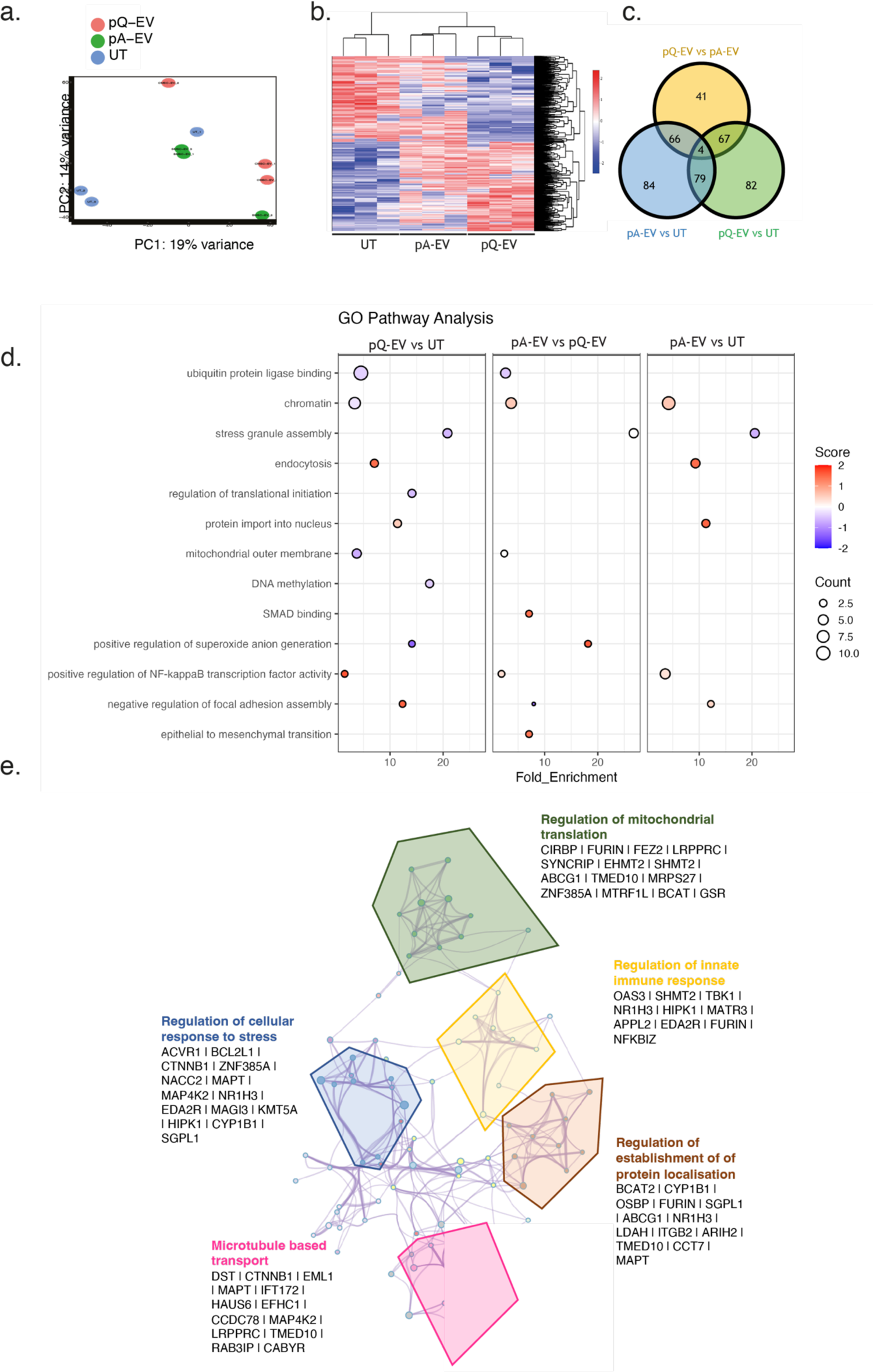
3D MCF7 cultures show significantly different RNA-seq profiles upon 72 hour pQ-EV vs pA-EV incubations 3D MCF7 cultures were untreated (UT) or incubated for 72 hours with pQ-EV/ pA-EV treatments before RNA extraction for NGS RNA sequencing. **a**.) Principle component analysis is shown, where each point represents 1 replicate, n=3. **b**.) Heatmap shows all hits, clustered through average linkage. **c**.) Venn Diagram shows distribution of all significant hits (p<0.001), across the different conditions. **d**.) Interesting hits from GO pathway enrichment shown. **e.**) Network diagrams illustrate pathways affected, where each node represents a significantly enriched term. Node size is proportional to the number of the contributing genes. Similar terms with a high degree of redundancy were clustered, as depicted, including a description and the genes involved. Map shows pA-EV treated spheroids’ pathways significantly upregulated compared to pQ-EV treated spheroids.

**SFigure 8.**
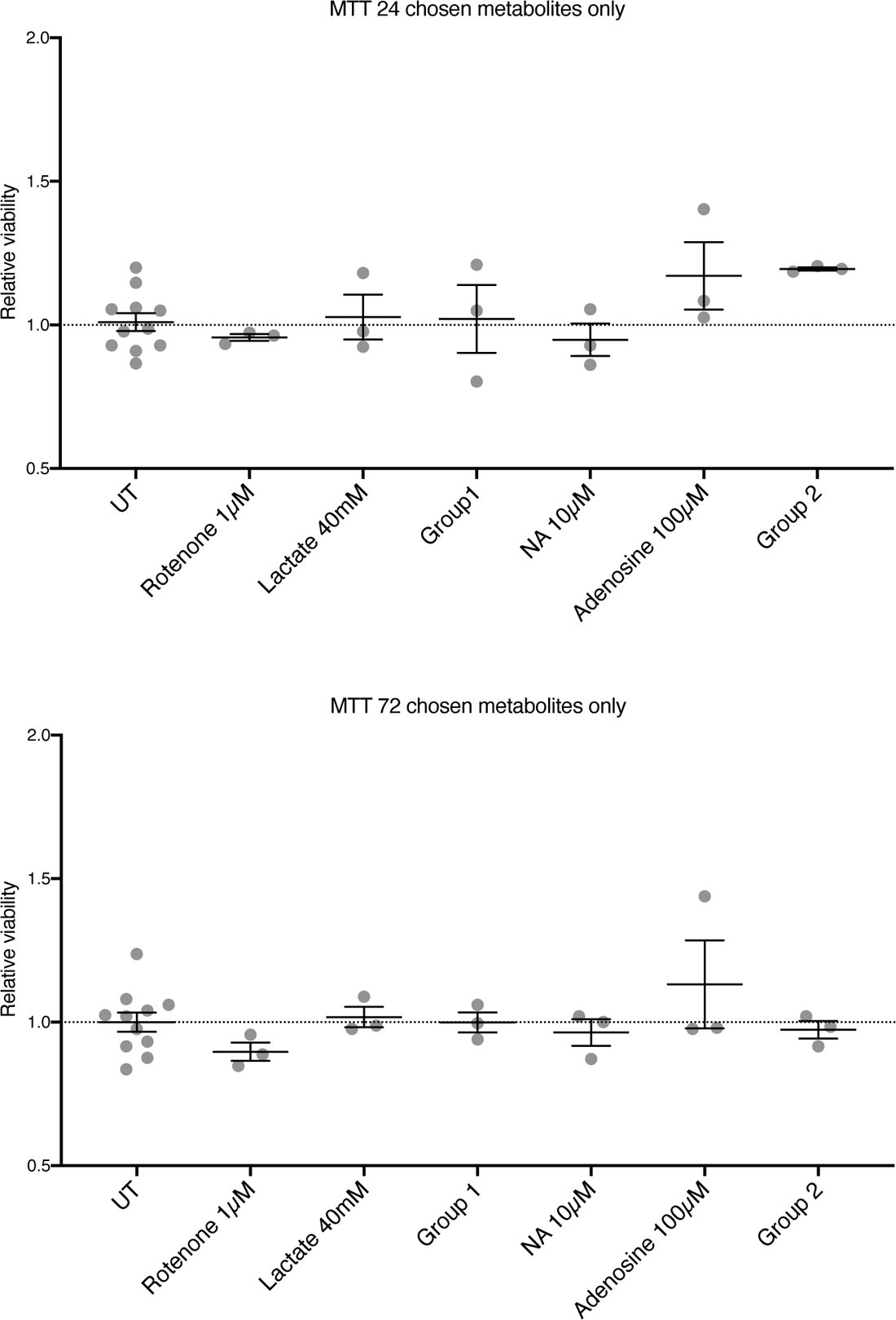
Metabolite MCF7 spheroid culture treatments do not significantly influence cell viability Cell viability was observed after MTT treatment upon 24/ 72 hours of MCF7 spheroid metabolite treatments. Graph represents mean absorbance normalised to UT cells ± EM. A one-way ANOVA with multiple comparisons showed no significant difference between samples, n=3.

**SFigure 9.**
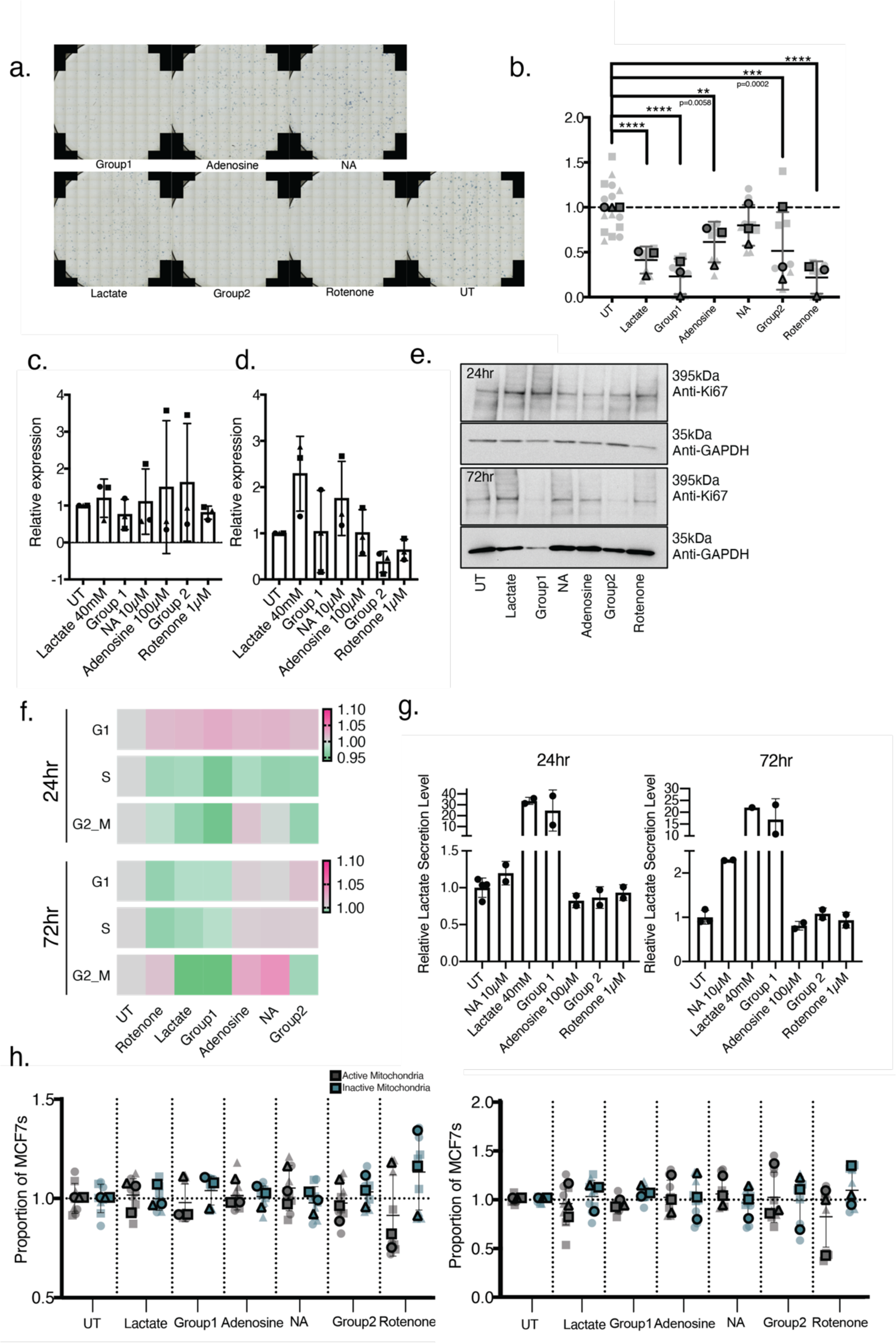
Optimisation of activity metabolites treatments. **a, b.**) A colony formation assay was used to assess the effect of the identified metabolite treatments over 2 weeks. MCF7s were seeded at a low density in 2D monolayer. These cultures were either untreated (UT) or treated with DMEM supplemented 40mM lactate, Group1 (40mM lactate+ 50mM glucose+ 8mM pyruvate), 10µM NA, 100µM Adenosine, Group2 (10µM NA+ 100µM Adenosine) or 1µM Rotenone treatments every 3 days for 2 weeks, after which colonies were fixed, stained and counted. Treatments were normalised to the UT control. **a**.) Representative images**. b.)** Mean colonies per condition ± SD plotted, and an unpaired t-test performed, n=3. **c-e.)** 3D MCF7 cultures were UT or incubated for 24/ 72 hours with metabolite treatments before lysates were collected for Ki67 expression Western blot analysis. Ki67 band density was normalised to a GAPDH loading control and then further normalised to UT samples to calculate a fold-change value. **c.)** Quantification of Ki67 expression after 24 hour treatment. **d.**) Quantification of Ki67 expression after 72 hour treatment **e.)** Representative blot shown from n=3 independent blots. **f.)** 3D MCF7 spheroids were UT or incubated for 24/72 hours metabolite treatments before being collected, dissociated with collagenase D, washed, fixed and stained with FxCycle™ PI/RNase Staining Solution (Life technologies; F10797). The stained single cell suspensions were subject to flow cytometry analysis. Graphs represent geometric mean fluorescent intensity per cell cycle phase, normalised to UT cells ± SD, with a minimum of 2000 cells analysed per 3 technical repeats. A 2-way ANOVA with multiple comparisons test was done, n=3. **g.)** 3D MCF7 lactate secretion levels were measured, providing an insight into the cellular levels of glycolytic metabolism upon respective metabolite treatments. Supernatants were collected upon each timepoint, and lactate levels quantified via the Lactate Glo^TM^ Assay (Promega, J5021). Graph depicts mean ± SEM concentration, n=2, with a minimum or 2 technical repeats run. **h.)** 3D MCF7 cultures were untreated (UT) or incubated for 24/72 hours with respective metabolite treatments and changes in mitochondrial respiration were evaluated via the JC-1 dye. Graph depicts mean ± SEM proportion of active and inactive mitochondria for each condition, normalised to UT cells. 3 technical repeats with a minimum of 2000 cells were measured per sample and an unpaired t-test completed between each condition, n=3.

## Notes

### Competing Interest Statement

The authors have declared no competing interest.

